# Loss of ADAMTS9 disrupts ciliogenesis and collagen homeostasis resulting in Nephronophthisis-like polycystic kidneys

**DOI:** 10.64898/2025.12.02.691883

**Authors:** Sydney Fischer, Karyn L. Robert, Manu Ahmed, Griffin I. Kane, Matthew A. Kavanaugh, Wei Wang, Pamela V. Tran, Prabhani U. Atukorale, Sumeda Nandadasa

## Abstract

*ADAMTS9* mutations cause the ciliopathies nephronophthisis and Joubert syndrome. Here we show that deletion of ADAMTS9 in the proximal nephron leads to polycystic kidney development in mice. In males, *Adamts9* deletion cause kidneys to become highly cystic but remain small without undergoing enlargement, causing early postnatal lethality. Female mice on the other hand, develop cystic kidneys but progress slowly. ADAMTS9 deletion disrupted ciliogenesis by the loss of ciliary transition zone (TZ) protein TMEM67 cleavage, leading to loss of the MKS/B9 module – a key component of the ciliary gate. Functional analysis of all eight ciliopathy patient variants of *ADAMTS9* identified to date, showed TMEM67 C-terminus failed to localize to the transition zone, thus disrupting a key regulatory mechanism in patient renal ciliogenesis. Modeling ADAMTS9-mediated TMEM67 cleavage utilizing our novel TMEM67-cleavage deficient mice revealed loss of TZ formation but not elevated canonical Wnt signaling as the underlying mechanism driving cystogenesis. We show that *Adamts9* deletion leads to comparatively intense interstitial collagen deposition, which likely restricts kidney enlargement resulting in the characteristically small kidney phenotype in nephronophthisis and increased immune response. By comparative analysis of four interconnected polycystic kidney models in addition to *Pkd1* and *Pkd2* deleted kidneys, we identify differential collagen homeostasis is the principle determining factor deciphering cystic kidney size and type.

## INTRODUCTION

Nephronophthisis (NPHP) is the most common genetic cause of end-stage renal failure in children and young adults and is caused by autosomal recessive mutations affecting the formation or function of renal primary cilia [1]. Key characteristics of NPHP include cyst formation in the corticomedullary region, high interstitial fibrosis, and disruption of the tubular basement membrane [2]. NPHP patients can also manifest a number of other ciliopathies, including JBTS (Joubert syndrome), which affects cerebellum formation; retinal abnormalities such as retinitis pigmentosa and coloboma; and a multitude of heart, liver and skeletal abnormalities [3, 4]. Unlike the more common polycystic kidney disease (PKD), which is caused by autosomal dominant or recessive mutations of the polycystin multi-protein complex genes [5], and characterized by highly enlarged polycystic kidneys, NPHP kidneys form smaller or normal-sized cystic kidneys. To date, NPHP is associated with over 20 genes, which are involved in the formation or the function of primary cilia [6]. While defective cilia function remains the only common denominator amongst different forms of cystic kidney disease, the underlying molecular mechanisms leading to cyst formation downstream of cilia function remain highly debated and elusive [7]. What specific factors determine cystic kidney size and sub-type by different ciliopathy variants is also unknown.

Until recently, the extracellular matrix metalloprotease ADAMTS9 (A Disintegrin and Metalloproteinase with Thrombospondin motifs, family member 9) was thought to be exclusively acting on extracellular matrix clearance by its ability to proteolytically cleave the chondroitin sulphate proteoglycans Versican and Aggrecan [8–13]. Unexpectedly, we discovered that homozygous recessive mutations of *ADAMTS9* result in the ciliopathies NPHP and JBTS and its loss of function disrupts ciliogenesis in human cells and mouse embryos [14, 15]. Since then, six additional *ADAMTS9* variants have been identified and a total of eight *ADAMTS9* variants are now known to cause NPHP and JBTS [14, 16, 17] (**Fig. 1A**). We also discovered that catalytically active ADAMTS9 localizes to the base of the cilium in Rab11 vesicles and its catalytic activity is essential for cilia formation [15]. However, the underlying molecular mechanism affecting ciliogenesis was unknown. To uncover its ciliary substrates, we conducted an advanced proteomics screen utilizing the iTRAQ-TAILS N-terminomics technique [18]. This identified TMEM67, a key component of the ciliary transition zone (TZ) as a novel substrate of ADAMTS9 [19, 20]. Characterizing ADAMTS9-mediated TMEM67 cleavage revealed that this cleavage event is required for the formation of the MKS/B9 module of the ciliary TZ by generating a TMEM67 C-terminal functional-form (TMEM67 Δ342) which translocates to the cilium [20]. The MKS/B9 module, in part forms the extracellular and transmembrane compartment of the TZ, also known as the transition one necklace [21]. Generation of a novel TMEM67-cleavage deficient mouse model and characterization of *TMEM67* ciliopathy patient variants of the cleavage site revealed, TMEM67-cleavage is essential for normal development of the mammalian kidney [20]. *Adamts9*-null mice on the other-hand die during gastrulation and ADAMTS9’s function during kidney development or NPHP progression was unknown [22]. Modeling ADAMTS9 loss in human kidney organoids further demonstrated the significance of *ADAMTS9* for normal kidney organoid development and cell signaling [16], however the role of ADAMTS9 in mammalian kidney development has remained uninvestigated.

**Figure 1.**
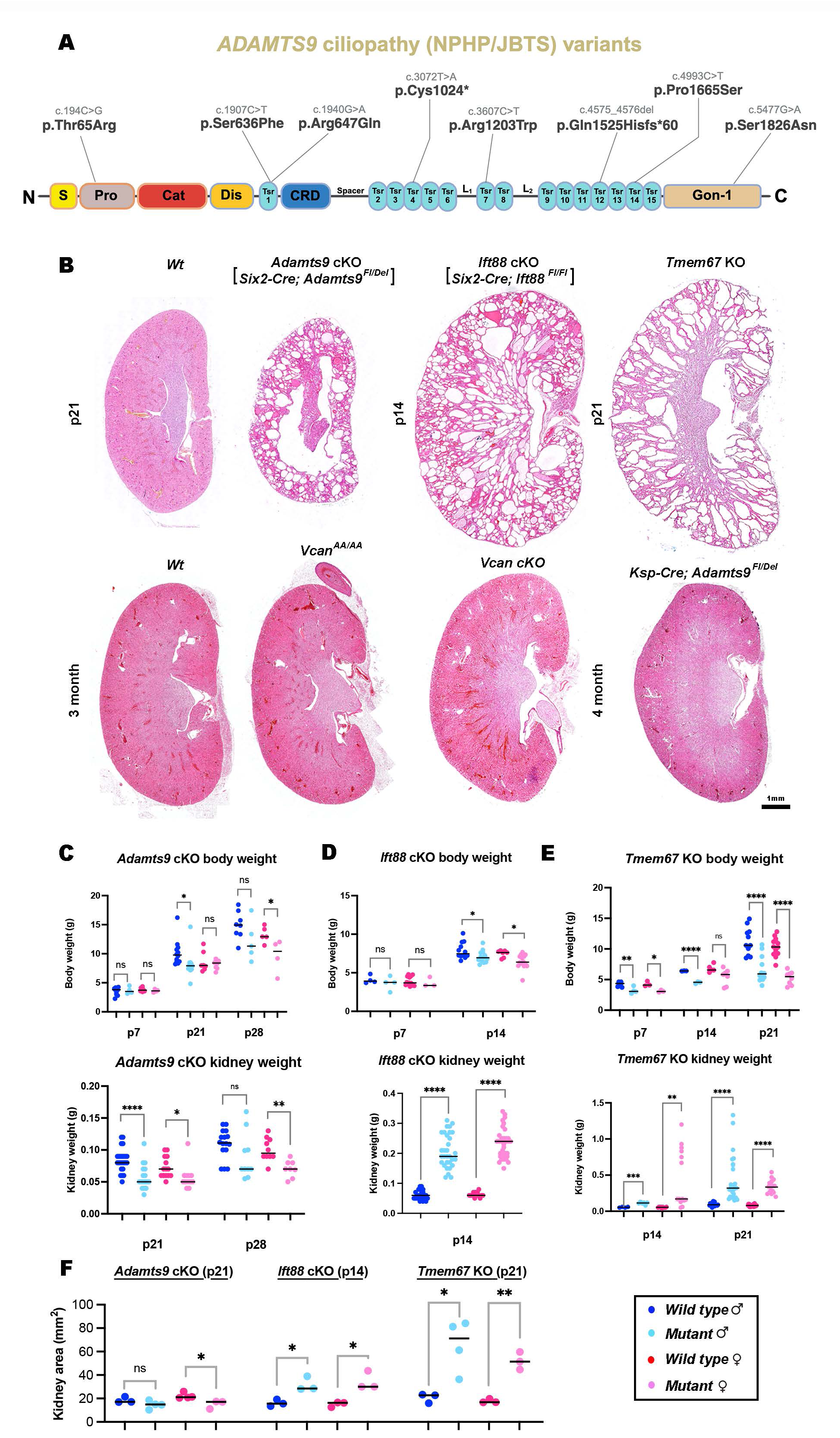
Deletion of *Adamts9* in the proximal nephron causes small but highly cystic kidneys in mice. (**A**) Ciliopathy patient variants of *ADAMTS9* mapped to its domain structures. S, signal peptide; Pro, Pro-domain; Cat, catalytic domain; Dis, Disintegrin domain; Tsr; Thrombospondin type-1 repeat; CRD, cysteine rich domain; L_1_, linker-1; L_2_, linker-2; Gon-1, Gon-1 domain. (**B**) Hematoxylin and eosin stained kidney sections of male kidneys (n= at least 3/ group). *Adamts9* conditional deletion in the proximal nephron results in highly cystic but relatively small polycystic kidneys in comparison to *Ift88* cKO and *Tmem67* KO kidneys. Mutation of the ADAMTS cleavage site of the versican GAG-β domain (*Vcan^AA/AA^* mice), or conditional deletion of versican in the mouse kidney (*Vcan* cKO), and conditional deletion of *Adamts9* in the ureteric lineage (collecting ducts), do not result in polycystic kidneys. Scale bar = 1 mm. (**C-E**) Body weights (top row) and kidney weights (bottom row) of male and female *Adamts9* cKO, *Ift88* cKO, *Tmem67* KO, and their control littermates show *Adamts9* cKO polycystic kidneys remain small in size even at their terminal stage, compared to other cystic kidney models and Wt kidneys. Error bars indicate ± SEM, **** indicates a p-value <0.0001, **<0.01, *<0.05 calculated by two-tailed unpaired *t*-test. For body weights, p=0.373 (p21 male *Adamts9* cKO vs Wt), p=0.0399 (p28 female *Adamts9* cKO vs Wt), p=0.0321 (p14 male *Ift88* cKO vs Wt), p=0.0102 (p14 female *Ift88* cKO vs Wt), p=0.0023 (p7 male *Tmem67* KO vs Wt), p=0.0106 (p7 female *Tmem67* KO vs Wt), and p<0.0001 (p14 male *Tmem67* KO vs Wt, p21 male *Tmem67* KO vs Wt*, and* p21 female *Tmem67* KO vs Wt). For kidney weights, p=0.0254 (p21 female *Adamts9* cKO vs Wt), p=0.0012 (p28 female *Adamts9* cKO vs Wt), p=0.0002 (p14 male *Adamts9* cKO vs Wt, p14 male *Ift88* cKO vs Wt, p14 female *Ift88* cKO vs Wt, p21 male *Tmem67* KO vs Wt, and p21 female *Tmem67* KO vs Wt). (**F**) Kidney area quantification at the indicated time points (n= at least 3/ sex/ genotype). *Ift88* cKO mice do not survive beyond two weeks and their kidney area was measured at p14. Error bars indicate Mean ± SEM. * indicates a *p*-value < 0.5 calculated by two-tailed unpaired *t*-test with p=0.0280 (female *Adamts9* cKO vs *Wt*), p=0.0158 (male *Ift88* cKO vs *Wt*), p=0.0145 (female *Ift88* cKO vs *Wt*), p=0.0190 (male *Tmem67* KO vs *Wt*), and p=0.0015 (female *Tmem67* KO vs *Wt*).

Here we conditionally deleted *Adamts9* in the murine kidney nephric mesenchyme lineage utilizing the proximal tubule and podocyte specific *Six2-*Cre and in the ureteric bud lineage utilizing the distal nephron-specific *Ksp*-Cre-drivers [23, 24]. We show that loss of ADAMTS9 in the proximal nephron (but not the distal nephron), mimics the human disease NPHP and results in small but highly cystic kidneys with high levels of interstitial collagen accumulation. We show that conditional deletion in males results in a more severe phenotype compared to female mice, and that ADAMTS9 is required for TMEM67 cleavage and transition zone assembly in renal primary cilia. In addition, we find that ADAMTS9 deletion leads to a complete loss of the MKS/B9 module of the TZ. Further characterizing TMEM67 cleavage in the murine kidney utilizing the novel TMEM67-cleavage deficient mouse model revealed key differences in canonical Wnt signaling in comparison to *Tmem67*-null and *Adamts9* deficient cystic kidneys, excluding elevated canonical Wnt signaling as the common driving force for renal cyst formation. Functional analysis of *ADAMTS9* ciliopathy variants highlighted the significance of ADAMTS9-mediated TMEM67 cleavage in disease etiology, revealing loss of TMEM67 C-terminus localization to the ciliary transition zone as a crucial regulatory step. We postulate that loss of ciliogenesis coupled with loss of collagen homeostasis unique to *Adamts9* deletion by its proteolytic cleavage of MT_1_-MMP (MMP14) leads to characteristically small but highly cystic kidneys, mimicking NPHP in mice.

## RESULTS

### ADAMTS9 deletion in the proximal nephron causes cystic kidney development

Homozygous, loss of function mutations in *ADAMTS9* causes nephronophthisis. To date, eight disease causing alleles have been reported in cystic kidney patients, whose mutations are spread throughout the different domains composing the large extracellular matrix metalloproteinase (**Fig. 1A**). However, the underlying molecular mechanism of cyst formation nor its function in mammalian kidney development has been investigated. To assess the impact of *Adamts9* deletion in the mouse kidney, we first assessed its expression pattern in the mouse nephron utilizing the Kidney Cell Explorer database, developed by comparing single cell RNA sequencing data (scRNAseq) from male and female mouse kidneys [25]. This revealed *Adamts9* expression was limited to the proximal end (podocytes, parietal epithelium and proximal tubules) in both male and female nephrons, but male mice showed significantly higher level of expression in the 3^rd^ segment of the proximal tubules in comparison to female kidneys (**Fig. S1A**). Strikingly, the novel ADAMTS9 substrate TMEM67, also showed a similar expression pattern and was expressed at high levels in the 3^rd^ segment of male proximal tubules (**Fig. S1A-B**). We also examined the expression of other versicanases belonging to the ADAMTS superfamily and *Vcan* in the kidney cell explorer. This showed no bias in expression comparing male and female kidneys (**Fig. S1C**). We therefore carried out *Adamts9* conditional deletion utilizing the *Six2*-Cre driver, which expresses Cre recombinase in the nephrogenic lineage [23], targeting the proximal nephron (here forth referred to as *Adamts9* cKO mice). Male *Adamts9* cKO mice showed highly cystic, small kidneys compared to *wild type (Wt)* and littermate control (heterozygous and *Adamts9^Fl/+^)* mouse kidneys of the same sex by 21 days of age (**Fig. 1B**). Strikingly, cystic kidney development in female *Adamts9* cKO mice progressed much more slowly and only showed highly cystic kidneys at 3 months of age (**Fig. S1D**). At the terminal stage of disease progression, *Adamts9* cKO male mice took on a hunched posture in their 3^rd^ week of life and were distinctly small with an abnormal gait compared to heterozygous males or cKO female littermates, who were indistinguishable from healthy female mice at 3 weeks of age (**Movie S1, Movie S2 and Fig. S2A**). We utilized conditional deletion of *Ift88* using the same Cre driver as a model completely lacking renal ciliogenesis and *Tmem67* KO mice as a model lacking the ADAMTS9 substrate of the ciliary transition zone, in comparison to *Adamts9* cKO kidneys. Deletion of *Ift88* results in a complete loss of cilia formation [26], while deletion of TMEM67 results in abnormally long or short cilia in the murine kidney [20]. Both these models developed highly cystic kidneys which were greatly enlarged compared to *Adamts9* cKO kidneys (**Fig. 1B-F**).

We previously developed a versican cleavage deficient mouse model (*Vcan^AA/AA^*) in which the ADAMTS9 cleavage site of the Versican GAG β-domain was mutated uncleavable [27]. Neither the *Vcan^AA/AA^* mice or conditional deletion of versican in the kidney [*Six2*-Cre; *Vcan^Fl/Fl^* (*Vcan cKO*), or *Ksp*-Cre; *Vcan^Fl/Fl^* (data not shown)] did not result in cystic kidneys (**Fig. 1B**). These data indicated cystic kidney development was unrelated to ADAMTS9’s versican GAG-β domain cleavage function in the murine kidney. This supported our previous findings in human RPE-1 cells [15], and also suggested versican and its cleavage are dispensable for normal kidney development. Conditional deletion of *Adamts9* in the ureteric lineage utilizing *Ksp*-Cre did not develop cystic kidneys, validating the Kidney Cell Explorer expression data, indicating ADAMTS9 function is highly specific to the proximal nephron (**Fig. 1B**). *Adamts9* cKO mice had lower body weights and kidney weights compared to littermate control mice in both sexes (**Fig. 1C**). In stark contrast, *Ift88* cKO mice and *Tmem67* KO mice had significantly lower body weights and significantly higher kidney weights resulting from their large polycystic kidneys (**Fig. 1D-E**). Quantification of kidney area also showed a similar result, where *Adamts9* cKO kidneys remained small while *Ift88* cKO and *Tmem67* KO kidneys were highly enlarged (**Fig. 1F**). Periodic Acid-Schiff (PAS) staining, a valuable tool to assess kidney injury showed proteinaceous, polysaccharide-rich nephric casts throughout the *Adamts9* cKO cystic kidneys (**Fig. S2B-C**). These data revealed, loss of ADAMTS9 in the proximal nephron of the murine kidney produces small but highly cystic kidneys, recapitulating the human disease NPHP, thereby providing a valuable platform to study the disease mechanism by which loss of *ADAMTS9* causes nephronophthisis.

### *Adamts9* cKO kidneys undergo normal nephrogenesis and forms cysts postnatally

To investigate when cyst formation occurred in *Adamts9* cKO mice, we harvested kidneys on the last day of embryonic development (E18.5) and at 7 days of age (p7) and analyzed their histopathology in midline sections. Both male and female *Adamts9* cKO kidneys did not show distinct cystic dilations by hematoxylin and eosin staining but were slightly smaller in size compared to *Wt* and the other cystic kidney models (**Fig. 2A**). *Ift88* cKO and *Tmem67* KO kidneys showed a large number of nephric tubules which were abnormally dilated at E18.5. High magnification analysis of *Adamts9* cKO kidney medullary regions showed large areas filled with mesenchymal cells in comparison to *Wt* kidneys (**Fig. 2B**). At p7 all three models showed cystic dilations and *Ift88* cKO kidneys in contrast to the others were considerably larger in size (**Fig. 2A**). To investigate if *Adamts9* cKO kidneys despite being highly cystic, remain smaller due to formation of fewer number of nephrons were formed in them during embryonic development, we first quantified the number of glomeruli observed per midline section in E18.5 and p7 kidneys (**Fig. 2C**). This analysis revealed no significant differences in the number of glomeruli present in any of the cystic kidney models at E18.5, but fewer glomeruli were present in *Ift88* cKO kidneys by p7. Secondly, staining of *Wt* and *Adamts9* cKO kidneys at E18.5 with the nephron segmental markers LTL (proximal tubules) and DBA (collecting duct) reveled comparable staining for both lineages (**Fig. 2D**). However, high magnification images showed the LTL staining of the *Adamts9* cKO kidney cortices were morphologically distinct from the *Wt* kidneys, with tubular dilations, presumably marking the beginning of cystic expansion, which was not apparent by hematoxylin and eosin staining. Staining of male *Wt* and *Ift88* cKO kidneys at E18.5 with LTL and DBA showed a similar result with distinctly dilated proximal tubules (**Fig. S3A**). Combined, these results suggest that ADAMTS9 enzymatic activity is required for normal kidney development in mice and its absence leads to postnatal cystic expansion.

**Figure 2.**
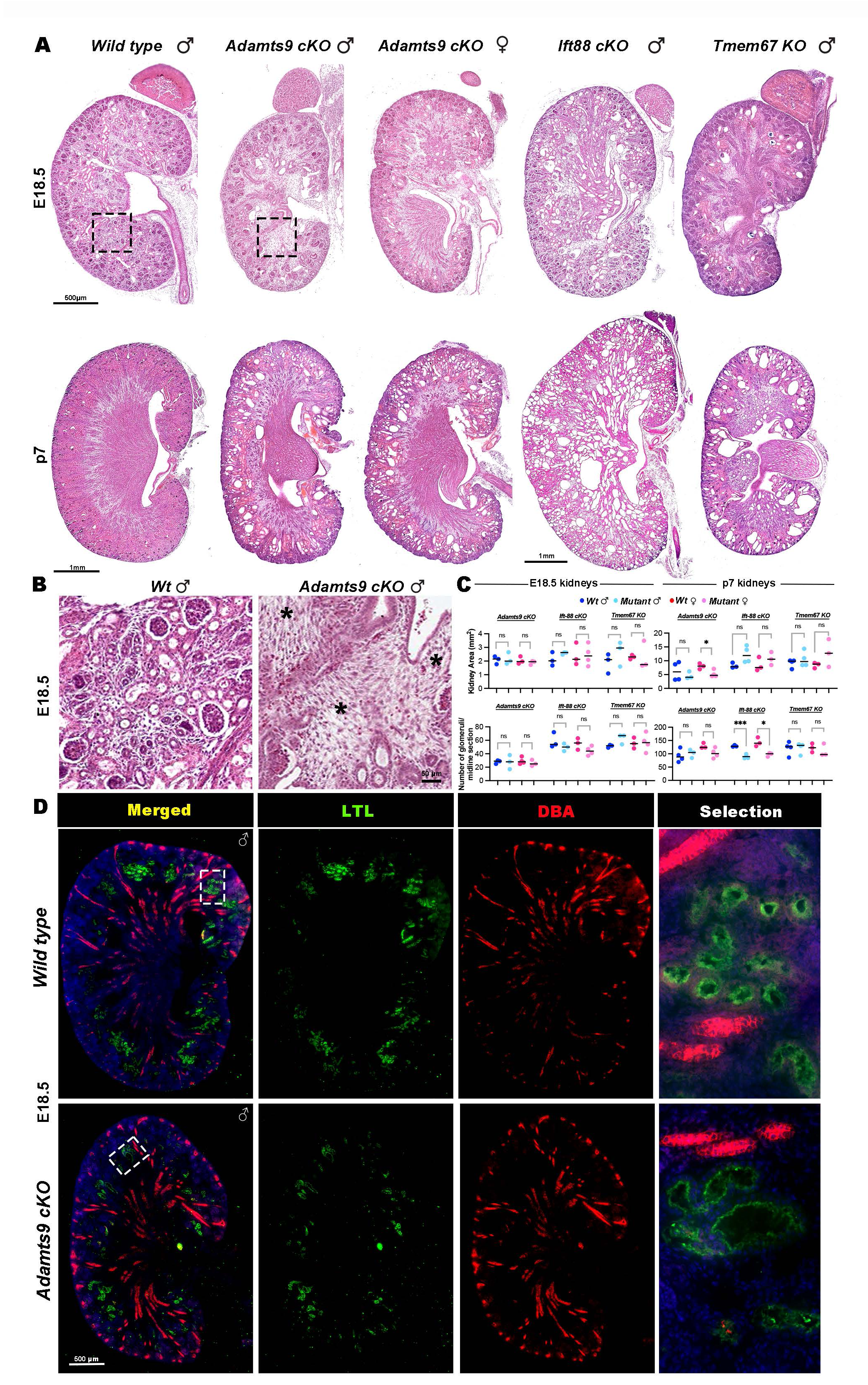
Normal embryonic nephrogenesis and postnatal cyst formation in *Adamts9 cKO* kidneys. (**A-B**) Hematoxylin and eosin staining of embryonic day 18 and a half (E18.5, top row) and post-natal day 7 (p7, bottom row) kidneys. Both male and female *Adamts9* cKO kidneys show normal renal histopathology at the end of embryonic development and show cystic expansion at p7 while *Ift88* cKO and *Tmem67* KO kidneys show onset of cysts at E18.5 (n=3-4 kidneys/sex/genotype). Boxed areas are shown in higher magnification in **B**. * indicate large areas with mesenchyme-like cells present in the *Adamts9* cKO kidney sections. (**C**) Quantification of kidney area and number of glomeruli in midline kidney sections of E18.5 (left) and p7 (right) hematoxylin and eosin stained kidney sections (n=3-4/sex/genotype). Error bars indicate Mean ± SEM. * indicates a *p*-value <0.5 and ***<0.001 calculated by two-tailed unpaired *t*-test. (**D**) Male, E18.5 Wt and *Adamts9* cKO kidneys stained for proximal tubules (LTL, green) and medullary regions, showing abnormally enlarged morphologies of *Adamts9* cKO proximal tubules (green) in comparison to Wt tubules. Scale bars in **A** are = 500 µm (E18.5 kidneys, top row) and 1 mm in p7 kidneys (bottom row), 50 µm in **B** and 500 µm in **D**.

### ADAMTS9 is required for ciliary necklace and renal primary cilia formation

To investigate ciliogenesis in *Adamts9* cKO kidneys, we first used freeze-fracture scanning electron microscopy (SEM), which revealed distinct primary cilia in the renal tubules of *Wt* kidneys but their near complete absence in the *Adamts9* cKO kidneys (**Fig. 3A**). Only very few cystic epithelial cells in *Adamts9* cKO kidneys possessed primary cilia, which were extremely short (red arrowheads). Additional analysis of cilia ultrastructure by transmission electron microscopy (TEM) showed a complete loss of the ciliary “necklace”, the distinct transmembrane and extracellular compartment of the TZ [21], in the short cilia observed in *Adamts9* cKO kidneys (**Fig. 3B**). We also observed a large number of partially formed cilia which had failed to extend a long axoneme beyond the TZ. To asses ciliogenesis in *Adamts9* cKO kidneys by immunostaining, we stained kidney cryo-sections with the ciliary membrane marker ARL13B and the basal body marker FOP (FGFR1 oncogene partner). Confirming the EM observations, immunostaining also revealed an overall complete loss of ciliogenesis with few extremely short primary cilia present in the *Adamts9* cKO kidneys, compared to the distinct cilia seen in *Wt* kidneys (**Fig. 3C**). To investigate if loss of ciliogenesis is due to loss of TZ assembly in the *Adamts9* cKO kidneys, we cultured 21-day-old kidneys and analyzed TZ formation in ice-cold methanol fixed primary cell cultures. We previously identified six molecules belonging to the MKS/B9 module of the ciliary TZ that were lost by loss of ADAMTS9 or TMEM67 in RPE-1 (retinal pigment epithelial) cells [20]. We investigated TZ localization of these six molecules in renal primary cilia by immunostaining *Wt* and *Adamts9* cKO kidney primary cultures (**Fig. 3D, Fig. S4A-F**). The TZ molecules TCTN1, TCTN2, TCTN3, TMEM237 and B9D2 showed significantly lower TZ localization in *Adamts9* cKO kidneys while CC2DA levels were not significantly affected (**Fig. 3D**). These results supported the kidney TEM observations, in which the TZ necklace, composed of the tectonic complex (TCTN1-3) and other transmembrane components of the TZ [21], was completely lost in the *Adamts9* cKO renal primary cilia, a phenotype observed previously in *ADAMTS9*-null RPE-1 cells [20].

**Figure 3.**
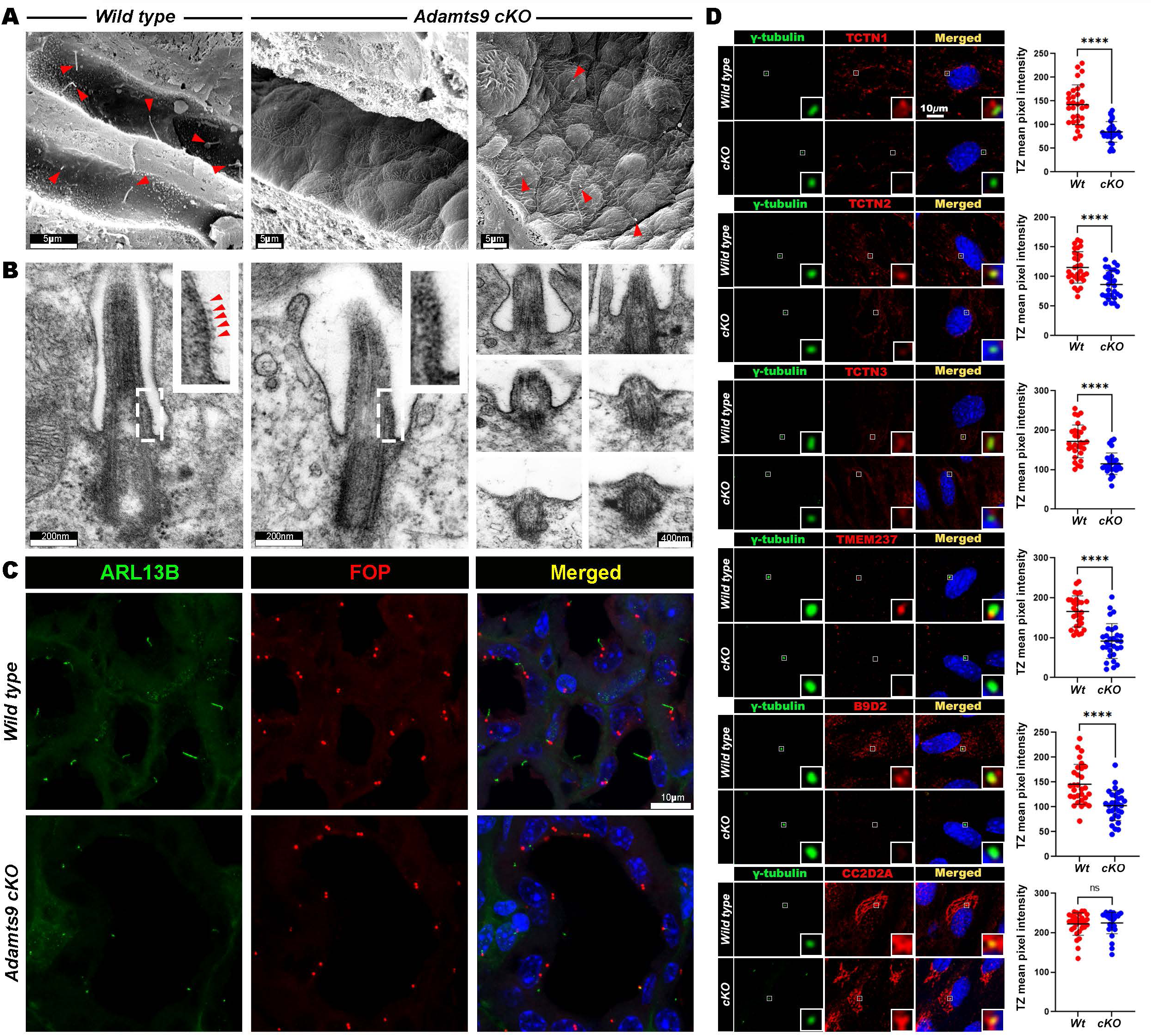
Loss of ciliogenesis in *Adamts9* cKO kidneys. (**A**) Freeze-fracture SEM imaging of male p21 *Wt* and *Adamts9* cKO kidneys (n=3) showing a complete loss of ciliogenesis or extremely short cilia (red arrow heads) present in *Adamts9* cKO cystic renal tubular epithelium compared to the long cilia seen in *Wild type*. (**B**) TEM imaging of renal primary cilia showing the transition zone “necklace” (red arrows) seen in *Wt* kidneys are absent in the short *Adamts9* cKO cilia while some mutant cilia were not extended beyond the cell surface. (**C**) Immunostaining of the ciliary axoneme marker, ARL13B (green), and the basal body marker, FOP (red), in p21 male *Wt* and *Adamts9* cKO kidneys showing loss of ciliation in the mutant cystic renal tubular epithelium. Inserts show high magnification of the boxed area (basal body/ TZ). (**D**) *Adamts9* cKO kidney primary cell cultures immunostained for the basal body (γ-tubulin, green) and the indicated transition zone markers (red) show a loss of MKS/B9 module formation in the *Adamts9* cKO cilia (n=3 experiments). Error bars indicate Mean ± SEM. **** indicates a *p*-value <0.0001 calculated by two-tailed unpaired *t*-test. Scale bars in **A** is 5 µm, 200 nm in **B,** 400 nm in **C**, and 10 µm in **D**.

### *Adamts9* is required for ECM homeostasis in the murine kidney

Since ADAMTS9 is also a *bonafide* extracellular matrix metalloproteinase, we investigated the overall impact of ADAMTS9 loss on ECM homeostasis in the murine kidney. Masson’s trichrome analysis showed large accumulations of collagen (blue staining) surrounding the glomeruli and in the interstitial matrix surrounding the cystic tubules in *Adamts9* cKO kidneys (**Fig. 4A**). Immunostaining for Collagen IV, which is limited to the glomeruli in *Wt* kidneys, showed high levels of staining throughout the *Adamts9* cKO cystic kidneys and in the glomerular basement membrane GBM (**Fig. 4B**). The loss of only a few other ECM associated proteins are known to cause polycystic kidney disease. The lamin-511 α subunit, Laminin α5 (*Lama5*) is the best characterized among these and has previously shown to cause polycystic kidneys when deleted in mice [28, 29]. Strikingly, immunostaining for Laminin α5 showed decreased, defused and non-fibrillar staining pattern in the GBM of *Adamts9* cKO kidneys compared to *Wt* kidneys (**Fig. 4C**). Further, ultrastructure analysis of the GBM by TEM showed thickened GBMs present in *Adamts9* cKO cystic kidneys (**Fig. 4D**). To our surprise, analysis of Laminin α5 and Collagen IV immunostaining in *Ift88* cKO cystic kidneys showed the opposite staining pattern (**Fig. S3B-D**). Laminin α5 staining was highly upregulated and Collagen IV staining was decreased in *Ift88* cKO glomeruli. These results indicate that while the GBM composition and morphology and overall ECM homeostasis is also severely impaired by loss of ADAMTS9 in the murine kidney, the underlying molecular mechanism driving these ECM dynamics may differ from that caused by loss of renal primary cilia alone (i.e., *Ift88* cKO).

**Figure 4.**
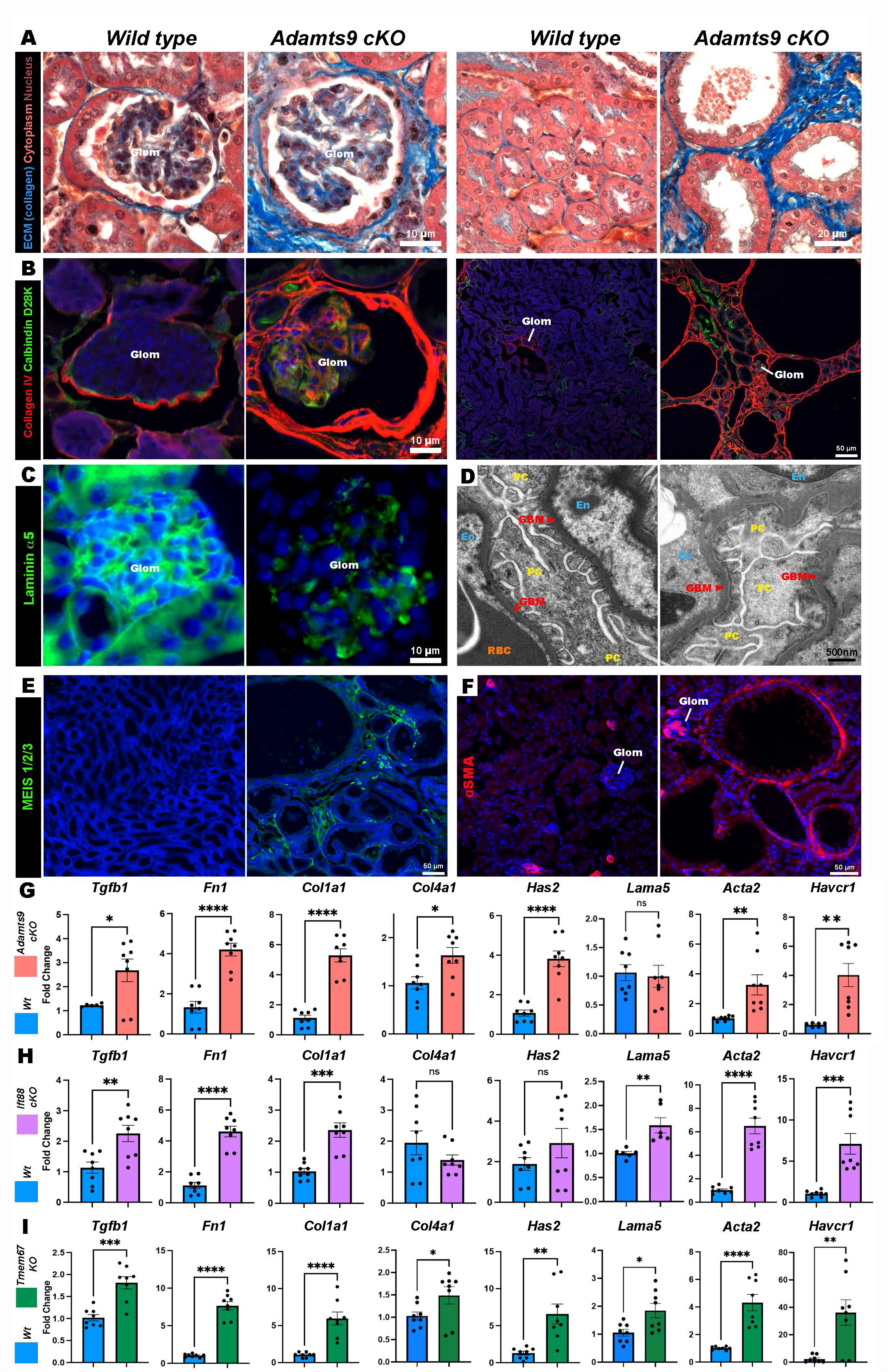
Adamts9 cKO cystic kidneys are severely impaired in extracellular matrix homeostasis. (A) Masson’s Trichrome staining of p21 male *Wt* and *Adamts9* cKO kidneys show collagen accumulation (blue) surrounding glomeruli (left) and cystic tubules (right). (B) Collagen IV (red) immunostaining which is limited to glomeruli in *Wt* kidneys, show intense staining surrounding the *Adamts9* cKO glomeruli (left) and throughout the kidney. Calbindin D28K immunostaining (green) which marks the distal convoluting tubules and collecting duct tubules show smaller tubules amongst larger cystic tubules in *Adamts9* cKO kidneys. (C) Laminin α5 ( green) staining seen in the *Wt* glomerular basement membrane (GBM) is decreased in *Adamts9* cKO glomeruli. (D) TEM images showing thickening of the glomerular basement membrane (GBM) of *Adamts9* cKO kidneys in comparison to *Wt* (n=3, each group). Podocytes (PC), endothelium (En), red blood cell (RBC). (**E-F**) Kidney sections stained with renal fibroblast marker MEIS 1/2/3 (green) (**E**) and the fibrosis marker α-SMA (red), (**F**), show *Adamts9* cKO kidneys are undergoing renal fibrosis. MEIS positive cells are absent in Wt kidneys and α-SMA is limited to vascular capillaries in Wt kidneys. (**G-I**) Quantitative, real time RT PCR (qRT-PCR) analysis of ECM-related genes (*Tgfb1, Fn1, Col1a1, Col4a1, Has2, Lama5*), the kidney injury (*Havcr1*), and fibrosis (*Acta2*) markers on cDNA synthesized from whole kidneys from male, p21 *Adamts9* cKO, *Tmem67* KO, and p14 *Ift88* cKO kidneys in comparison to their respective *Wt* littermates show altered gene regulation in the cystic kidneys (n=4 kidneys/group). Error bars indicate Mean ± SEM, **** indicates a p-value <0.0001,***<0.001,**<0.01,*<0.05 calculated by two-tailed unpaired *t*-test. Scale bars in **A** are 10 µm and 20 µm, 10 µm and 50 µm in **B**, 10 µm in **C**, 500 nm in **D**, 50 µm in **E** and **F**.

Immunostaining for the homeobox protein Meis1, which marks renal myofibroblasts [30, 31], showed accumulation of renal myofibroblasts throughout the interstitial ECM surrounding the cystic kidney tubules in the *Adamts9* cKO kidneys (**Fig. 4E**). Similarly, alpha smooth muscle actin (αSMA) staining, which is limited to the vasculature in *Wt* kidneys, also showed staining throughout the *Adamts9* cKO and *Ift88* cKO kidneys surrounding the cystic renal tubular epithelial cells and glomeruli (**Fig. 4F, Fig. S3E**). Together, these results indicate that *Adamts9* cKO cystic kidneys undergo fibrosis and have increased ECM deposition. Since *Adamts9* cKO and *Ift88* cKO kidneys showed different staining patterns, we inquired if the ECM dynamics are resulted from deregulated cell signaling and transcriptional changes to ECM-related genes. Quantitative, real-time RT-PCR (qRT-PCR) analysis for TGF-β (*Tgfb1*), which is a master regulator of ECM related gene transcription and fibrosis [32], showed significantly upregulated levels in the *Adamts9* cKO kidneys. Similarly, transcript levels of the ECM molecules fibronectin (*Fn1*), collagens type I and IV (*Col1a*, *Col4a*) and hyaluronan synthase 2 (*Has2*) showed highly upregulated levels in the *Adamts9* cKO cystic kidneys (**Fig. 4G**). Conversely, laminin α5 which showed decreased staining in the *Adamts9* cKO kidneys, showed decreased transcript levels (*Lama5*) although was not significant. The renal fibrosis indicator smooth muscle alpha2 actin (*Acta2*) and the kidney injury marker, Kim1 (*Havcr1*) also showed significantly high transcript levels in the *Adamts9* cKO kidneys (**Fig. 4G**). Cystic kidneys from *Ift88* cKO or *Tmem67* KO mice also showed similar deregulations of these ECM transcripts, indicating that loss of cilia function can indeed regulate ECM dynamics by deregulated gene transcription in cystic kidneys (**Fig. 4H-I**). Intriguingly, *Ift88* cKO kidneys, which showed highly increased Laminin α5 staining in the GBM and a moderate decrease in Col-IV staining, showed correlating gene transcription (**Fig. 4H)**. *Lama5* transcription was upregulated while *Col4a1* was downregulated in *Ift88* cKO kidneys. In *Tmem67* KO kidneys, all ECM-related genes analyzed were upregulated (**Fig. 4I)**. Combined, these data reveal the presence of highly complex etiologies in different cystic kidney models leading to abnormal ECM homeostasis, which may be governed in part by transcriptional changes. Since loss of Versican nor mutating its known ADAMTS9-cleavage site in the GAG-β domain did not lead to cystic kidney development and were indistinguishable from *Wt* mouse kidneys even upon aging (**Fig. 1B**), we did not further investigate versican or its proteolysis in this study.

### ADAMTS9-mediated TMEM67 cleavage in the murine kidney is required for normal kidney development

Utilizing the iTRAQ-TAILS (isobaric tagging for relative and absolute quantitation, combined with terminal amine isotopic labeling of substrates) technique [18], henceforth referred to as N-terminomics for simplicity, we recently identified and validated the TZ molecule, TMEM67, as a novel substrate of ADAMTS9 [20]. ADAMTS9-mediated TMEM67 extracellular domain cleavage resulted in shedding of the TMEM67 cysteine rich domain (CRD) and translocation of the cleaved TMEM67 C-terminal half to the TZ for ciliogenesis (**Fig. 5A**). To investigate if ADAMTS9 is required for TMEM67 cleavage in the murine kidney, we performed western blot analysis of *Wt* and *Adamts9* cKO kidney primary cultures. This revealed significantly reduced TMEM67 N-terminal fragment shedding in the *Adamts9* cKO culture medium, indicating *Adamts9* cKO kidneys are deficient for TMEM67 cleavage (**Fig. 5B**). To study TMEM67 cleavage directly, we mutated the ADAMTS9 cleavage site of TMEM67 and generated a TMEM67 cleavage deficient mouse model (*Tmem67^ΔCLE^* mice) [20]. Similar to the *Tmem67*-null, *Tmem67^ΔCLE/ΔCLE^* mice also developed large cystic kidneys by 14 days of age (**Fig. 5C**). Both male and female *Tmem67^ΔCLE/ΔCLE^* mice had significantly reduced body weights and significantly increased kidney weights and kidney areas (**Fig. 5D**). Analysis of TMEM67 cleavage comparing *Wt* and *Tmem67^ΔCLE/ΔCLE^* renal primary cultures revealed lack of TMEM67 cleavage in *Tmem67^ΔCLE/ΔCLE^* kidneys (**Fig. S5A**). PAS staining in *Tmem67^ΔCLE/ΔCLE^ and Tmem67-*null cystic kidneys showed proteinaceous, polysaccharide-rich nephric casts throughout the cystic kidneys (**Fig. S5B**). Similar to *Tmem67*-null (**Fig. 2A**), *Tmem67^ΔCLE/ΔCLE^* mice also showed cystic dilations at E18.5 (**Fig. S5C-D**). Staining of E18.5 *Tmem67^ΔCLE/ΔCLE^* kidneys with DBA and LTL lectins showed overall comparable staining levels to that of *Wt* kidneys, but distinctly larger LTL labeled tubules in the cortex (**Fig. S5E**). We previously showed that *Tmem67^ΔCLE/ΔCLE^* kidneys had reduced ciliogenesis and formed both significantly long and short renal cilia by EM analysis, and both these cilia types lacked formation of the TZ necklace [20]. Immunostaining for the ciliary membrane marker ARL13B and the basal body marker FOP confirmed these observations and showed abnormal ciliogenesis in both *Tmem67-*null and *Tmem67^ΔCLE/ΔCLE^* kidneys (**Fig. 5E**). Intriguingly, the abnormally long primary cilia in the *Tmem67* mutant kidneys showed very low ARL13B intensities while the extremely short cilia showed intense ARL13B levels (**Fig. 5F**), indicating ARL13B and membrane trafficking dynamics may be severely impaired in these abnormal cilia. We also observed significant nuclear staining of ARL13B in the *Tmem67* mutant cystic kidneys (**Fig. 5E,G**). Interestingly, the ARL13B antibody used in these assays targets the unique C-terminus of ARL13B, when GFP tagged has been previously shown to localize to the nucleus [33]. Intriguingly, this ARL13B nuclear localization was not observed in the *Adamts9* cKO cystic kidney model, which suggested that the dynamic regulation of the ARL13B C-terminus may be unique to the TMEM67 loss-of-function cystic epithelium. Combined, these data reveal ADAMTS9 mediates TMEM67 cleavage in the murine kidney and is required to prevent polycystic kidney development in mice, but *Tmem67^ΔCLE/ΔCLE^* kidneys more closely modeled *Tmem67*-null kidneys, rather than *Adamts9* conditional deletion in mice.

**Figure 5.**
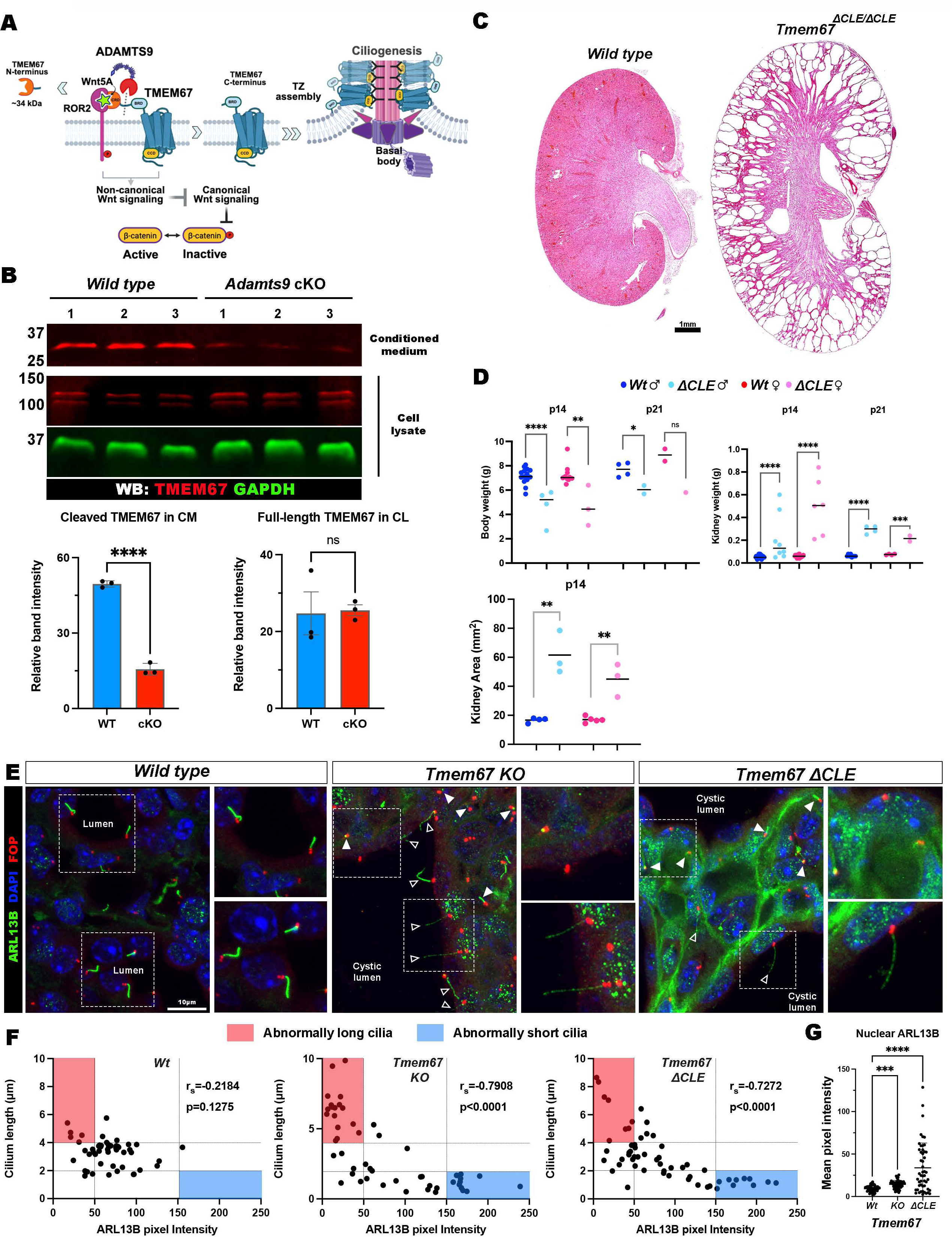
ADAMTS9 is essential for TMEM67 cleavage and renal ciliogenesis in the kidney. (A) Cartoon summarizing ADAMTS9 cleavage of the TMEM67 extracellular domain and the identification of two functional forms of TMEM67. The full-length form present at the cell surface is essential for transducing non-canonical Wnt signaling with ROR2, while the ADAMTS9-cleaved TMEM67 C-terminus translocate to the ciliary TZ, and is required for ciliogenesis. ADAMTS9-mediated TMEM67 cleavage releases an extracellular 34kDa N-terminal fragment. (B) Western blot analysis of kidney primary cultures assaying for TMEM67 cleavage show loss of the N-terminal 34 kDaTMEM67 fragment in the conditioned medium of *Adamts9* cKO cultures and its presence in *Wt* kidneys (n=3 cultures). The full-length TMEM67 protein level in the cell layer only showed a slight increase in comparison. TMEM67 (red) and GAPDH (green). (C) Mutating the ADAMTS9 cleavage site of TMEM67 to be uncleavable in *Tmem67^ΔCLE/ΔCLE^* mice leads to highly cystic kidneys. Hematoxylin and eosin staining of p14 Male mice are shown (n=3 each group). (D) Body weight, kidney weight, and kidney area quantifications of p14 and p21 *Wt* and *Tmem67^ΔCLE/ΔCLE^* mice from each sex. Note most *Tmem67^ΔCLE/ΔCLE^* mice are lethal embryonically and mice that are born rarely survives past day 14, leading to limited data of p21 kidneys. (E) Immunostaining of *Wt*, *Tmem67* KO and *ΔCLE* kidneys with the ciliary axoneme marker, ARL13b (green), and the basal body marker, FOP (red), show extremely short (white arrowheads) and abnormally long (open white arrowheads), renal cilia present in the cystic lumens of both *Tmem67* KO and *ΔCLE* kidneys in comparison to the *Wt* kidney tubules. (F) Quantification of cilia length and ARL13B pixel intensity correlation show abnormally long cilia with low ARL13B intensities (red box) and abnormally short cilia with intense ARL13B staining (blue box) in *Tmem67* KO and *ΔCLE* kidneys compared to *Wt* kidneys (n=3 kidneys, 30 cilia/ genotype). Statistical significance was determined by a two-tailed non-parameteric Spearman correlation (*Wt* -r_s_ = -0.2184, p=0.1275; *Tmem67* KO -r_s_ = -0.7908, p<0.0001; *Tmem67 ΔCLE* -r_s_ = -0.7272, p<0.0001). (G) Nuclear localized ARL13B quantification show increased staining in *Tmem67* KO and *ΔCLE* kidneys (n=3 kidneys, 30 nuclei/ genotype). **** indicates a *p*-value <0.0001 and ***<0.001in two-way ANOVA + Tukey’s multiple comparison test for statistical significance. Scale bar in **C** is 1 mm and 10 µm in **E**.

### Defective transition zone assembly in TMEM67 cleavage deficient kidneys

We previously showed that ADAMTS9-mediated TMEM67 cleavage is required for normal TZ assembly in RPE-1 cells and that *Tmem67^ΔCLE/ΔCLE^* renal primary cilia lacked the TZ necklace formation by TEM analysis [20]. To investigate renal primary cilia TZ assembly further, we established primary cultures from *Tmem67^ΔCLE/ΔCLE^* and *Tmem67* KO kidneys and compared them to *Wt* kidney cultures from their corresponding littermates (**Fig. 6A-F, Fig. S6A-F**). Intriguingly, similar to *Adamts9* cKO renal primary cultures, 5 TZ molecules (TCTN1-3, TMEM237 and B9D2) showed significantly decreased TZ localization while CC2D2A TZ localization was unaffected (**Fig. 6D**). This also suggested that CC2D2A stability at the TZ may be differentially regulated or differentially dependent on TMEM67 levels when comparing renal primary cilia and RPE-1 cells. Combined, these results show that loss of ADAMTS9, TMEM67 or its cleavage, severely impaired normal renal primary cilia formation and disrupted the formation of the TZ MKS/B9 module which forms the TZ necklace. Compared to *Adamts9* cKO kidneys, *Tmem67* mutants having both extremely long and short cilia suggest that ADAMTS9 may have more ciliary substrates which have not been identified yet, leading to a more severe loss of ciliation in the cystic renal tubular epithelium.

**Figure 6.**
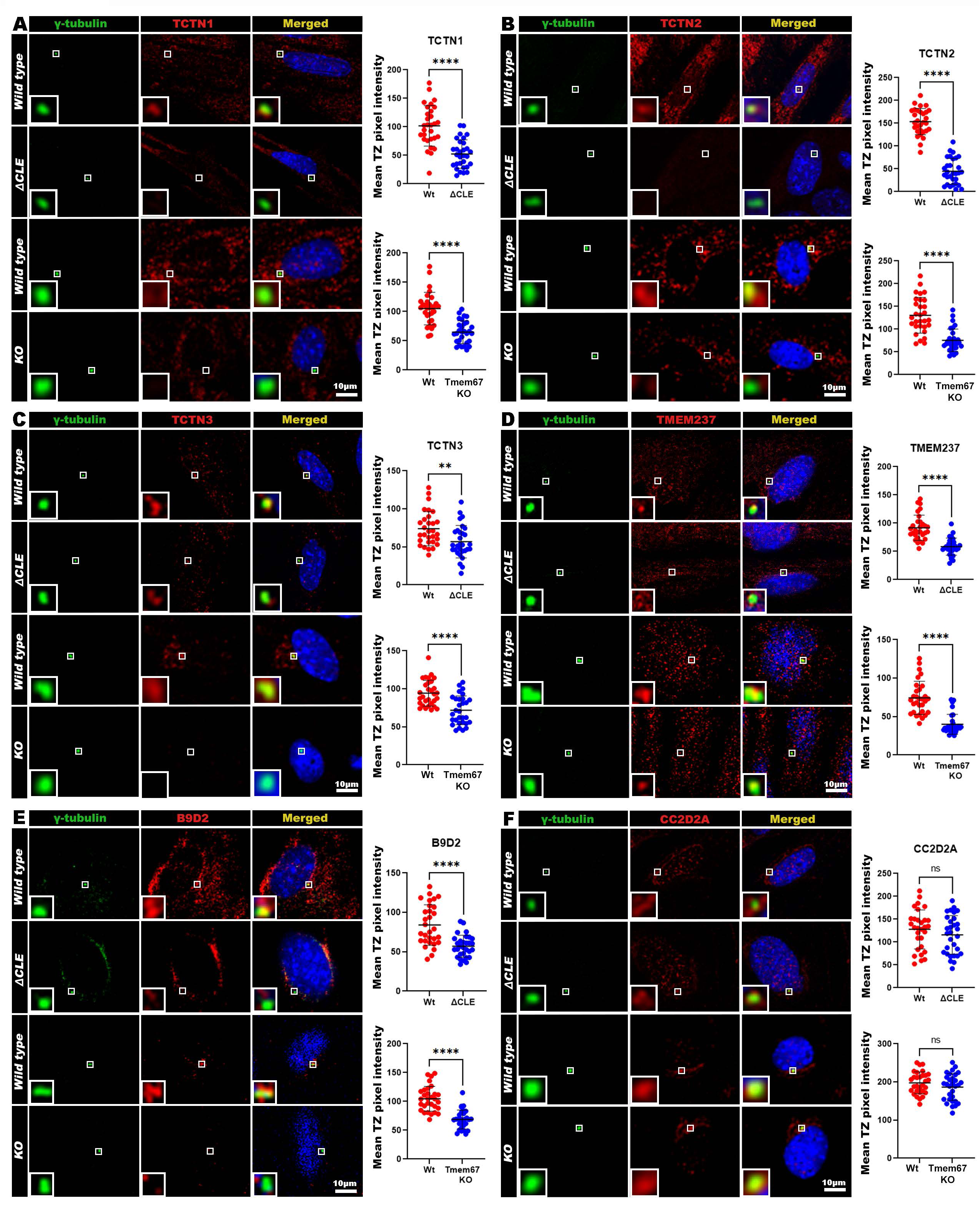
Los of MKS/B9 module proteins of the renal primary cilia of TMEM67 cleavage deficient and TMEM67-null kidneys. (**A-F**) Serum starved and ice cold methanol fixed *Wt, Tmem67^ΔCLE/ΔCLE^*, and *Tmem67* KO primary kidney cultures immunostained for the basal body marker, γ-tubulin (green) and the transition zone MKS/B9 module markers (red), show significantly lower staining for TCTN1-3, TMEM237 and B9D2 in the mutant TZs while CC2D2A is unaffected. Inserts indicate high magnification images of the transition zone of each cell (n=3 experiments, 30 TZs per group). Error bars indicate Mean ± SEM, **** indicates a *p*-value <0.0001 and **<0.01 calculated by two-tailed unpaired *t*-test. Scale bars in **A-F** are 10 µm.

### Defective ECM homeostasis in TMEM67 cleavage deficient cystic kidneys

To investigate if loss of TMEM67 cleavage can result in ECM homeostasis defects, we analyzed the ECM signature of *Tmem67^ΔCLE/ΔCLE^* kidneys. TEM analysis of glomeruli showed thicker GBM present in the *Tmem67^ΔCLE/ΔCLE^* kidneys (**Fig. 7A**). Masson’s trichrome staining confirmed these observations and showed increased levels of collagen deposited surrounding the glomeruli and also in the interstitial ECM surrounding the cystic tubules of *Tmem67^ΔCLE/ΔCLE^* kidneys (**Fig. 7B**). Similar to *Adamts9* cKO cystic kidneys, but dissimilar to *Ift88* cKO kidneys, Collagen-IV staining showed increased staining in the glomeruli and surrounding the cystic tubules of both *Tmem67 KO* and *11CLE* kidneys (**Fig. 7C**). Strikingly, Laminin α5 staining showed intense staining in the *Tmem67 KO* and *11CLE* glomeruli similar to *Ift88* cKO but dissimilar to *Adamts9* cKO kidneys which show decreased staining (**Fig. 7D**). α-SMA staining showed robust staining throughout both *Tmem67 KO* and *11CLE* kidneys (**Fig. 7E)**. qRT-PCR analysis for renal fibrosis and ECM markers showed a similar upregulation seen to that of *Tmem67-*null kidneys, which indicated these molecular changes were occurring by deregulated gene transcription and cell signaling events downstream of defective cilia formation (**Fig. 7F**). Of note, consistent with the immunostaining data, *Lama5* transcript levels were upregulated in both *Tmem67 KO* and *11CLE* kidneys and in *Ift88* cKO kidneys in contrast to *Adamts9* cKO kidneys, which showed decreased *Lama5* transcription. Intriguingly, in ADPKD, Laminin α5 expression is highly upregulated and has been shown to promote proliferation of the cystic renal epithelial cells [34, 35]. Thus the increased Laminin α5 levels in *Tmem67* mutant kidneys and *Ift88* cKO kidneys indicate greater similarity to the molecular signature of ADPKD cystic kidneys than to that of *Adamts9* cKO kidneys.

**Figure 7.**
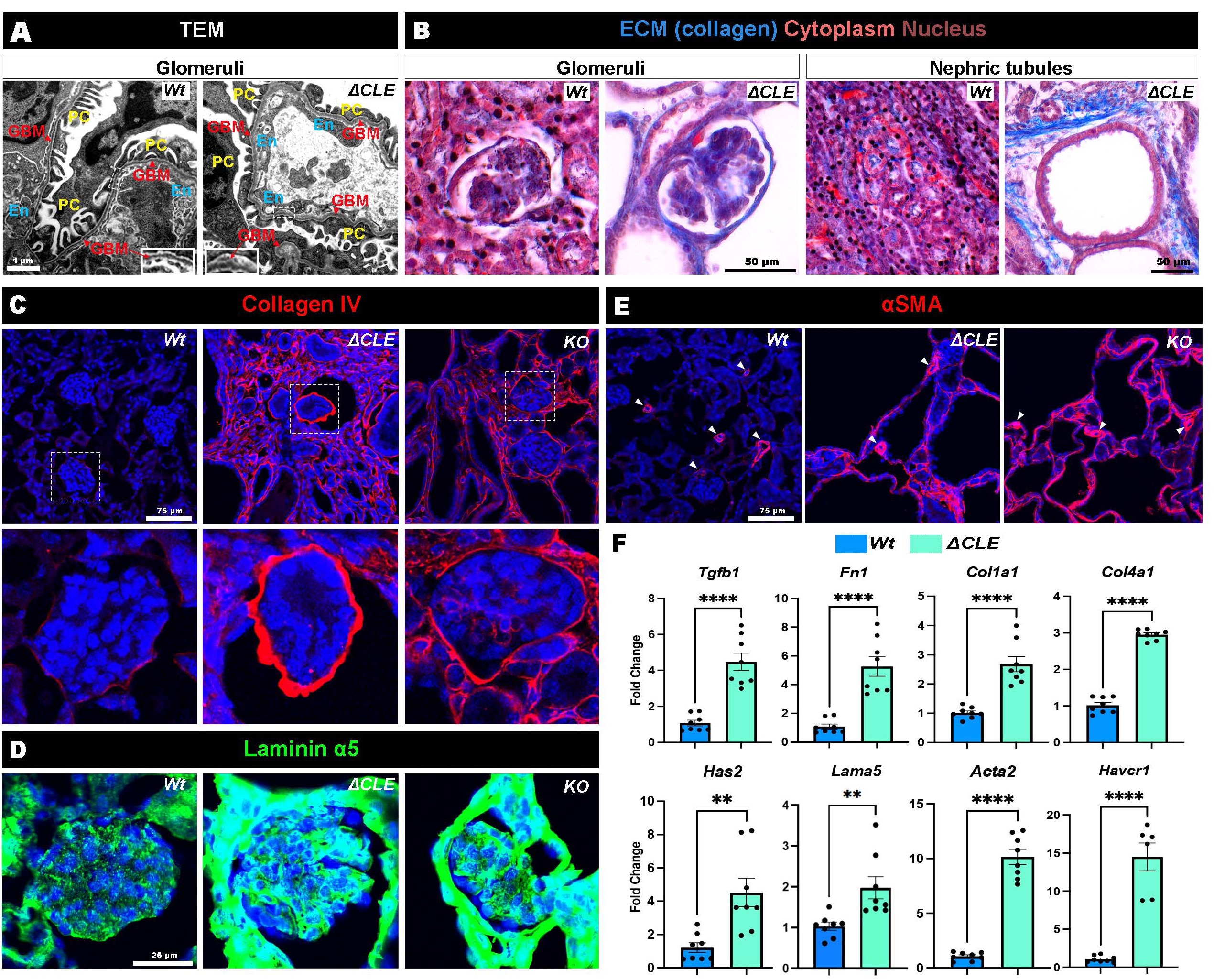
ECM homeostasis defects in TMEM67 cleavage deficient and TMEM67-null kidneys. (A) TEM images of p14 male *Wt* and *Tmem67^ΔCLE/ΔCLE^* kidneys show thickening of the glomerular basement membrane (GBM, red arrows). Boxed areas are shown in high magnification in the inserts. PC, podocytes; En, endothelium. (n=3 kidneys/ genotype) (B) Masson’s trichrome staining of p14 male *Wt* and *Tmem67 ΔCLE* kidneys show increased collagen (blue) deposition surrounding glomeruli and cystic tubules in the mutant kidneys (n=3 male kidneys/ genotype). (C) Immunostaining for Collagen IV (red) show increased staining throughout the kidneys of *Tmem67 ΔCLE* and *Tmem67* KO cystic kidneys (n=3 male kidneys/ group). Boxed area indicates a glomerulus shown in high magnification. (D) Intense laminin α5 staining (green) surrounding the glomeruli of *Tmem67 ΔCLE* and *Tmem67* KO cystic kidneys (n=3 male kidneys/ group). (E) α-SMA staining (red) which is limited to the vascular capillaries (arrow heads) in *Wt* kidneys, show intense staining throughout *Tmem67 ΔCLE* and *Tmem67* KO cystic kidneys (n=3 male kidneys/ group). (F) Quantitative, real time RT PCR (qRT-PCR) analysis of ECM-related genes (*Tgfb1, Fn1, Col1a1, Col4a1, Has2, Lama5*), kidney injury (*Havcr1*), and fibrosis (*Acta2*) markers of p14 male *Wt* and *Tmem67^ΔCLE/ΔCLE^*kidneys show increased transcription (n=4 kidneys/ genotype). Error bars indicate Mean ± SEM, **** indicates a p-value <0.0001,***<0.001, and **<0.01 calculated by two-tailed unpaired *t*-test. Scale bar in **A** is 1 µm, 50 µm in **B**, 75 µm in **C** and **E**, and 25 µm in **D**.

### Differential regulation of canonical Wnt signaling in *Adamts9* cKO and *Tmem67^ΔCLE^* kidneys

To investigate what signaling pathways maybe differentially regulating these differential gene expression patterns observed amongst the interconnected cystic kidney models, we investigated the Wnt signaling pathway in the four cystic models. Balanced Wnt singling is crucial for normal kidney development and disruption of both canonical and non-canonical Wnt signaling pathways have been associated with cystic kidney disease [36, 37]. More importantly to this study, in addition to its role in ciliary TZ assembly, TMEM67 is also involved in Wnt signal transduction by its N-terminal CRD, which forms a frizzled-like receptor for transducing non-canonical Wnt signaling mediated through Wnt5a, by forming a complex with ROR2 [38–42]. We recently showed that TMEM67 cleavage by ADAMTS9 mediates the balance between the full-length form present at the cell surface, which functions as a Wnt signaling receptor, and the cleaved C-terminal half which translocates to the TZ for its cilium assembly function [20] (summarized in **Fig. 5A**). In *Tmem67* KO mouse brains and embryonic fibroblasts (MEFs), non-canonical Wnt singling is lost and canonical Wnt signaling is highly upregulated. In contrast, *Tmem67 11CLE* mouse brains and MEFs, which forms a stable uncleavable full-length protein at the cell surface, transduces normal Wnt signaling [20].

To investigate the Wnt signaling signatures of the cystic kidneys, we first immunostained *Adamts9* cKO kidneys with the active β-catenin antibody, which specifically recognizes non-phosphorylated β-catenin, as a readout for canonical Wnt signaling (**Fig. 8A,E**). The *Adamts9* cKO kidneys showed highly elevated active β-catenin staining in the renal epithelial cells lining the cystic dilations while very low level of staining was observed in the healthy *Wt* littermate kidney tubules. *Ift88* cKO kidneys showed a similar pattern with increased active β-catenin staining in the renal epithelial cells (**Fig. 8B,E**). Strikingly, a similar increase in active β-catenin staining was only seen in the *Tmem67* KO kidneys and not in the *Tmem67 11CLE* kidneys, despite being highly cystic (**Fig. 8C-E**). These observations phenocopied our previous observations analyzing ependymal cells of the mouse brains of these mice and Wnt signaling assays conducted in MEFs, identifying canonical Wnt signaling as a major difference between the *Tmem67* KO and the *Tmem67 11CLE* mouse models [20]. To investigate non-canonical Wnt signaling signatures in the cystic kidneys, we investigated the expression of *Plod2, Lcor* and *Hadh* which are known downstream targets of the non-canonical Wnt signaling pathway, which we also previously observed to be differentially regulated in *Tmem67* KO and *11CLE* MEFs [20]. *Adamts9* and *Ift88* cKO kidneys showed no distinct regulatory pattern for these markers while *Tmem67* KO kidneys showed significantly low levels of *Plod2* and *Hadh* transcription (**Fig. 8F**). The *Tmem67 11CLE* kidneys on the other hand, which has constitutively cell surface localized full-length TMEM67 on their cell surfaces, showed highly elevated *Plod2, Lcor* and *Hadh* transcript levels (**Fig. 8F**). These results suggest that while both canonical and non-canonical Wnt signaling pathways are disrupted in the cystic renal tubular epithelium, increased canonical Wnt signaling alone, may not be sufficient to drive cystic kidney formation, since *Tmem6711CLE* kidneys are highly cystic, yet do not show increased canonical Wnt signaling (**Fig. 8D**).

**Figure 8.**
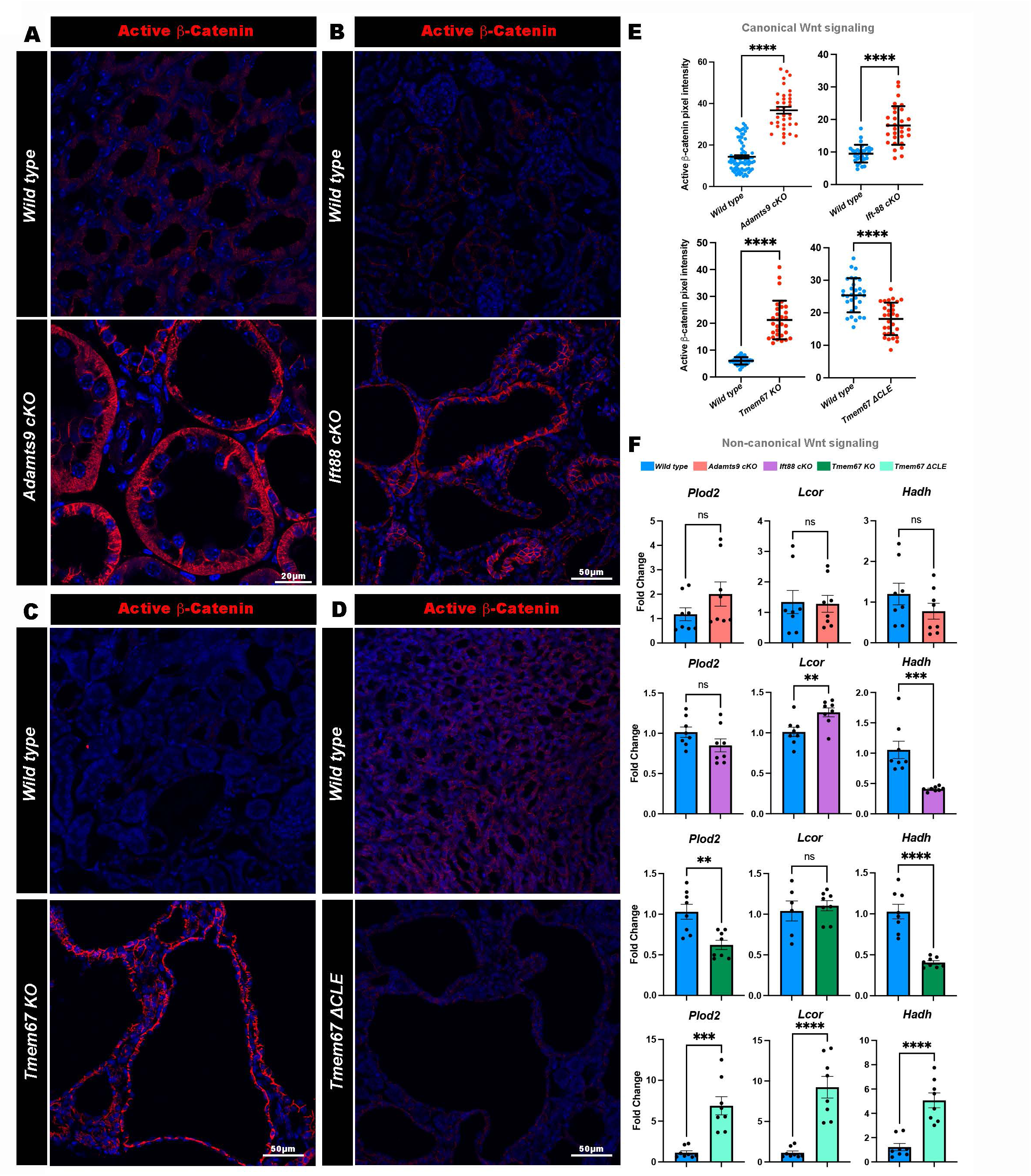
Differential canonical and non-canonical Wnt signaling modulation in polycystic kidney models. (**A-E**) Immunostaining for active β-catenin (red) at the terminal stage of male cystic kidneys and their respective *Wt* littermate kidneys show increased canonical Wnt signaling in *Adamts9* cKO (**A**), *Ift88* cKO (**B**) and *Tmem67* KO (**C**) polycystic kidneys but not in the *Tmem67 ΔCLE* kidneys (**D**), (n=3 kidneys/ genotype). (**E**) Quantification of active β-catenin pixel intensities (n=3 kidneys/ 30 tubules/ genotype). Error bars indicate Mean ± SEM. **** indicates a *p*-value < 0.0001 calculated by two-tailed unpaired *t*-test. (**F**) qRT-PCR analysis of known non-canonical Wnt signaling target genes *Plod2, Lcor* and *Hadh* in male *Adamts9* cKO, *Ift88* cKO*, Tmem67* KO, and *Tmem67 ΔCLE* kidneys in comparison to their respective *Wt* littermate kidneys show highly elevated non-canonical Wnt signaling in *Tmem67 ΔCLE* kidneys while *Tmem67* KO kidneys show reduced expression for *Plod2* and *Hadh* but not *Lcor* (n=4 kidneys/ genotype). Error bars indicate Mean ± SEM, **** indicates a p-value <0.0001, ***<0.001, and **<0.01 calculated by two-tailed unpaired *t*-test. Scale bar in **A** is 20 µm and 50 µm in **B-D**.

### Differential recruitment of macrophages and upregulation of inflammatory chemokine signaling in *Adamts9* cKO and other cystic kidneys

Inflammation and macrophage accumulation within the cystic kidney microenvironment is known to contribute to disease burden [43–45]. Remarkably, inhibition of inflammation has also been shown to slow down cystic kidney disease progression [46, 47]. We therefore analyzed the immune microenvironment in ADAMTS9 or TMEM67 deficient cystic kidney models in comparison to *Wt* littermates and *Ift88* cKO cystic kidneys which lacked primary cilia, and inquired if their immune responses were differentially regulated. This analysis was driven by our observations during freeze-fracture SEM, where we observed a large number of macrophages colonizing the cystic lumens of *Adamts9* cKO kidneys (**Fig. S7A**), which were also evident during TEM analysis (**Fig. S7B**), and by the fact that we did not observe similar large colonies of macrophages present in the lumen of any other cystic kidney models investigated in this study [(**Fig. S7C**) and [20]]. To probe this further, we investigated the immune signatures of all four cystic kidney models by flow cytometry analysis, in comparison to their *Wt* littermates in both sexes (**Fig. S7D-F**). Remarkably, despite having the smallest cystic kidney size, the *Adamts9* cKO kidneys showed the highest level of immune activity compared to *Ift88* cKO or *Tmem67* deficient cystic kidneys. Overall, there were high percentages of CD45+ immune cells, F4/80+ macrophages and CD11C+ dendritic or macrophage cells in all four cystic kidney models investigated compared to their littermates. Both male and female cystic kidneys showed increased levels comparable to one another and did not show any sex specific bias in the cystic kidney immune activation (**Fig. S7D-F**). To investigate chemokine signaling known to stimulate macrophage recruitment, we utilized qRT-PCR to test the expression level of MCP-1 (Monocyte chemoattractant protein-1, CCL2) [48]. *Adamts9* cKO kidneys showed ∼20 fold increase in MCP-1 (*Ccl2*) expression compared to *Wt* littermate kidneys. While *Ift88* cKO and *Tmem67* mutant cystic kidney models showed significantly increased levels of MCP-1, the upregulations were comparatively modest and less prominent to that seen in *Adamts9* cKO kidneys (**Fig. S7G**). CCR2 which acts as the primary receptor for MCP-1 [48], was transcriptionally upregulated in *Adamts9* and *Ift88* cKO kidneys, but downregulated in both *Tmem67* deficient kidneys (**Fig. S7G**). CCR4, another chemokine receptor, primarily found on T cells [49], showed a similar trend and showed upregulation in *Adamts9* and *Ift88* cKO kidneys but were either downregulated or not affected in the *Tmem67* deficient cystic kidney models (**Fig. S7G**). TNF-α (Tumor Necrosis Factor alpha), which is a master regulator of inflammation and immune cell activity [50], was highly upregulated in all four cystic kidney models with *Ift88* cKO kidneys showing the highest level of upregulation. These results show, in general, a highly immunogenic and inflamed kidney microenvironment present in all four cystic kidney models, but with differences in expression of key molecules involved in the immune responses of each model. The proinflammatory cytokines are produced by interstitial fibroblasts, inflammatory cells, and tubular epithelial cells in cystic kidneys [51]. The high levels of ECM present in the *Adamts9* cKO kidneys may be sequestering these cytokines, contributing to increased immunogenic activity and in the attachment of macrophages in the lumen of renal cysts, which were absent in others.

### Loss of TMEM67 C-terminus from the ciliary TZ is a common disease etiology in *ADAMTS9* ciliopathy patients

Polycystic kidney diagnosis is common to all known *ADAMTS9* ciliopathy patients identified to date. We therefore investigated whether these patient variants affected ciliogenesis and whether loss of TMEM67 cleavage is part of their disease etiology. To investigate the patient variant functionality in ciliogenesis, we generated all 8 variants (**Fig. 1A**) using site-directed mutagenesis and conducted rescue experiments in comparison to *Wild type* (WT), full-length human ADAMTS9 expression in *ADAMTS9*-null RPE-1 cells. Loss of ADAMTS9 resulted in loss of ciliogenesis and cilium length in RPE-1 cells, which was fully rescued by the introduction of WT ADAMTS9 but not the patient variants (**Fig. 9A**). Only the S1826N variant, which occurs in the C-terminus of the protein (in the unique Gon-1 domain), partially increased the percentage of ciliated cells and resulted in cilia lengths that were comparable to WT ADAMTS9 expression (**Fig. 9B,C**).

**Figure 9.**
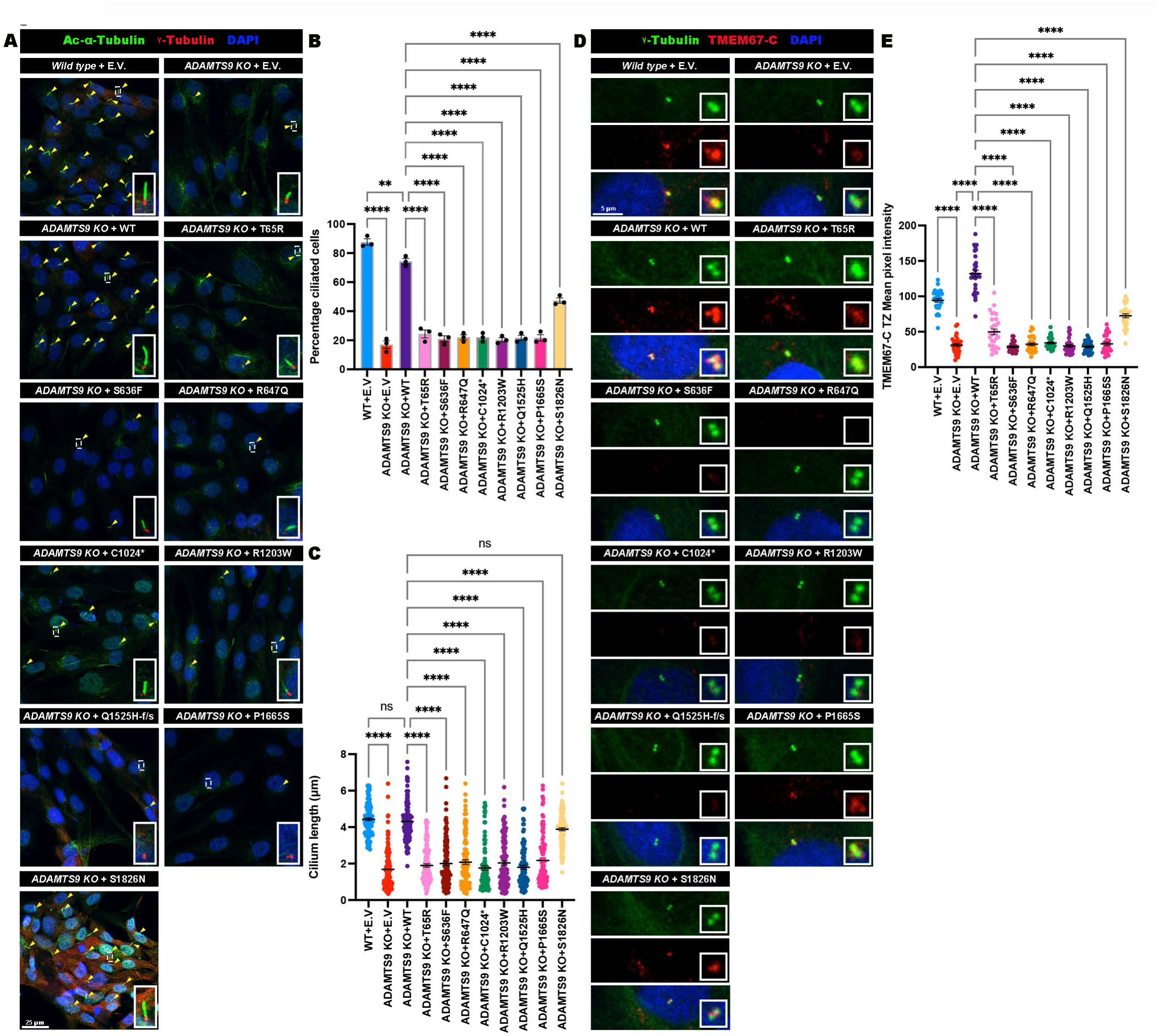
Ciliopathy patient variants of *ADAMTS9* are defective in ciliogenesis and transition zone localization of the TMEM67 C-terminus. (**A-C**) Immunostaining for primary cilia (yellow arrow heads) by Ac-α-Tubulin (green) and γ-tubulin (red) staining (**A**), in serum starved *Wt* or *ADAMTS9* KO RPE-1 cells transfected with empty vector (E.V.), WT, T65R, S636F, R647Q, C1024*, R1203W, Q1525H-f/s, P1665S and S1826N ADAMTS9 variants show significantly lower ciliogenesis (**B**), and cilia lengths (**C**) in comparison to WT ADAMTS9 transfection (n=3 experiments, 100 cells or 30 cilia/ group). Error bars indicate Mean ± SEM,**** indicates a *p*-value <0.0001 and **<0.01 in two-way ANOVA. (**D-E**) Immunostaining for TMEM67-C terminus (red) and γ-Tubulin (green) in ice-cold methanol fixed, serum starved cells (**D**), show significantly reduced transition zone localized TMEM67-C terminus in *ADAMTS9* KO RPE-1 cells, and cells transfected with ciliopathy variants (**E**), (n= 3 experiments, 30 transition zones/ group). Error bars indicate Mean ± SEM, **** indicates a *p*-value <0.0001 in two-way ANOVA. Scale bar is 25 µm in **A**, and 5 µm in **D**.

We previously demonstrated that *ADAMTS9*-null RPE-I cells lose the TMEM67 C-terminal half generated by its cleavage, localizing to the ciliary transition zone and introduction of the TMEM67 C-terminal half (TMEM67 Δ342) alone can partially restore cilium assembly and TZ assembly [20]. Utilizing an antibody generated against the TMEM67 C-terminus (TMEM67-C) [39], and ice-cold methanol fixation, we investigated the TMEM67 C-terminal localization status in *ADAMTS9* patient variants (**Fig. 9D, Fig. S8**). All variants had reduced localization of the TMEM67 C-terminus to the TZ in comparison to WT ADAMTS9 (**Fig. 9E**). These results indicate that TMEM67 cleavage may be at the center of a common disease mechanism affecting ciliogenesis in *ADAMTS9* ciliopathy patients leading to loss of ciliary function and NPHP kidney pathogenesis.

## Discussion

Here we demonstrate that loss of ADAMTS9 in the mouse kidney recapitulates the human cystic kidney disease NPHP, and ADAMTS9-mediated TMEM67 cleavage is essential for normal kidney development. We utilized four different cystic kidney models related to TMEM67 cleavage or loss of ciliogenesis (**Fig. 10A**), to investigate the similarities and dissimilarities in their molecular signatures, in comparison to their phenotypic manifestations (**Fig. 10B**). Our findings highlight that *Adamts9* conditional deletion in the nephrogenic lineage specifically causes small polycystic kidneys with high interstitial ECM accumulation, resembling the hallmarks of the human disease NPHP. In contrast, *Tmem67* loss of function mutants or *Ift88* nephrogenic lineage conditional deletion in mice leads to highly enlarged cystic kidneys, which are more reminiscent of an “ADPKD like” phenotype (**Fig. 10B**). While the four cystic kidney models in this study shared some common phenotypic manifestations such as cystogenesis, renal fibrosis, defective gene transcription and immune response, there were marked differences in their Wnt signaling signatures, their cilia phenotypes and their ECM levels (**Fig. 10B**). While both *Adamts9* and *Ift88* conditional deletion result in a complete loss of the cilium formation in the renal tubular epithelium, both *Tmem67* mutant models had a significant amount of ciliated cells, albeit they were either severely short or long.

**Figure 10.**
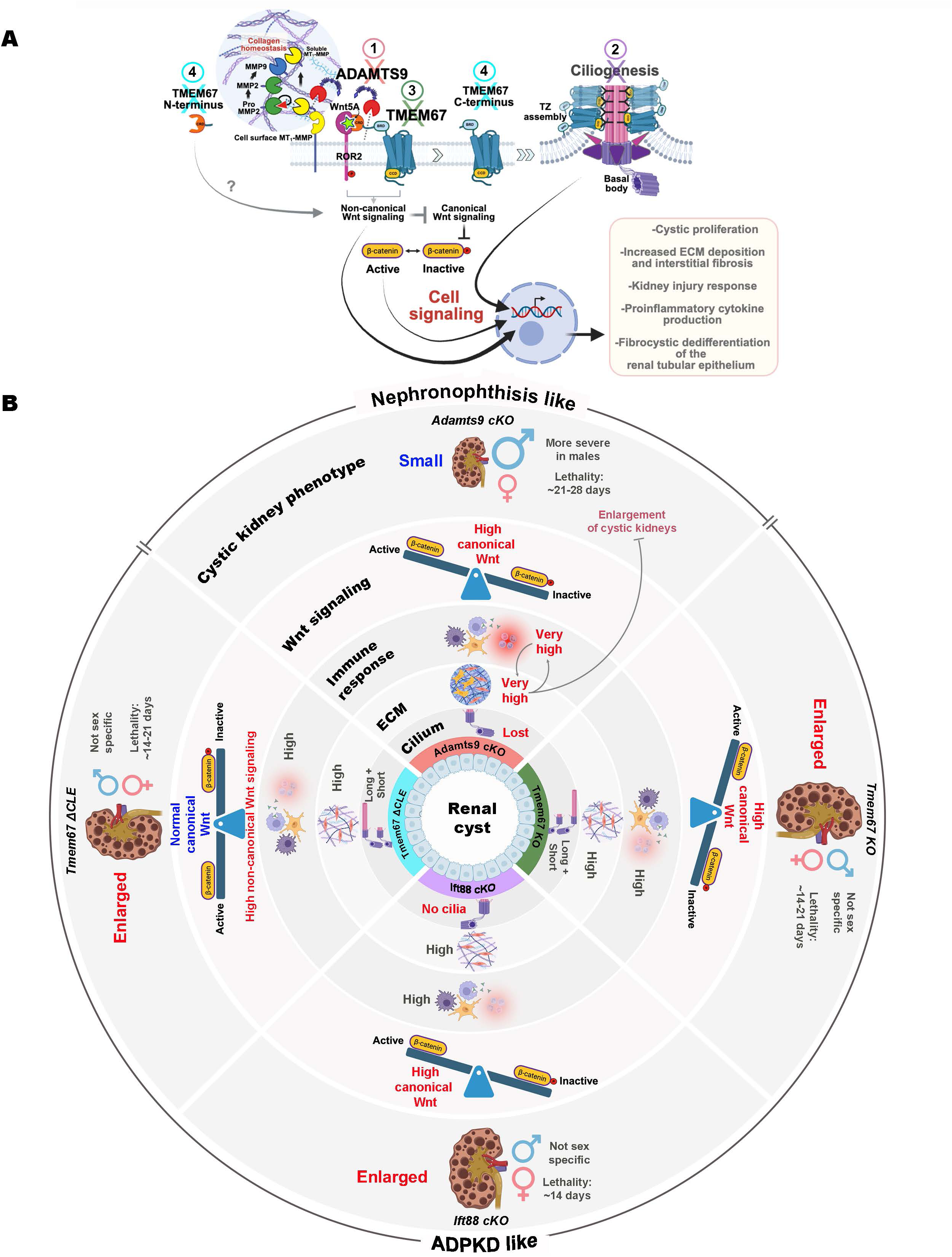
ADAMTS9 loss in the murine kidney leads to small NPHP-like kidneys while loss of IFT88 or TMEM67 function leads to large ADPKD like kidneys with or without defective canonical Wnt signaling. (A) Graphical summary depicting the four polycystic kidney models investigated and their point of impact in the pathway leading to loss of ciliogenesis and defective cell signaling. **1**- *Adamts9* cKO (red) impairs ECM homeostasis and TMEM67 extracellular domain cleavage, causing loss of cleaved TMEM67 C-terminus localizing to the TZ. **2**- *Ift88* cKO (purple) impairs cilium formation. **3**- *Tmem67*-null kidneys are deficient in Wnt signaling and TZ assembly. **4-** *Tmem67 ΔCLE* kidneys are deficient in forming the cleaved TMEM67 C- and N-terminal fragments, and are defective for TZ assembly and show elevated non-canonical Wnt signaling but normal canonical Wnt signaling. (B) Graphical summary depicting the key differences of the four cystic kidney models investigated and their phenotypic outcome causing either large ADPKD like kidneys (*Ift88* cKO, *Tmem67* KO and *ΔCLE* kidneys) or small NPHP like kidneys in the *Adamts9* cKO. *Adamts9* cKO in mice show onset of lethality in male mice ∼21-28 days and show a complete loss of ciliogenesis with very high levels of interstitial ECM, immune response and elevated canonical Wnt signaling. The ADPKD like cystic kidney models do not show a sex specificity with both males and female lethality onset ∼14-21 days and have relatively less ECM accumulation in comparison to *Adamts9* cKO kidneys. They have either no cilia (*Ift88* cKO) or extremely short or long cilia (*Tmem67* KO and *ΔCLE*) with or without elevated canonical Wnt signaling signatures and modestly increased immune responses. We postulate the elevated level of ECM surrounding the cystic tubules in *Adamts9* cKO cystic kidneys may be functioning as a stiff ECM scaffold preventing their enlargement, while also functioning as an ECM reservoir full of inflammatory cytokines and signaling molecules triggering macrophage attraction and a higher immune response.

Strikingly, both male and female *Adamts9* cKO kidneys have very high level of intestinal collagen, dramatically distinguishing them amongst the four cystic kidney models investigated in this study (**Fig. S9A-E**). While the other cystic kidney models do have somewhat elevated ECM related gene transcription and collagen deposition compared to their *Wt* littermates, the amount of collagen is distinctly lower compared to *Adamts9* cKO kidneys. How loss of function mutations related to cilia formation or cilia signaling can cause highly enlarged polycystic kidneys such as in ADPKD or small cystic kidneys such as in nephronophthisis has been a great unknown. This work sheds valuable insight into the role of renal fibrosis and interstitial ECM levels in the cystic kidneys may be an important deciphering factor of ADPKD-like vs NPHP phenotypic outcomes. To test this hypothesis directly, we analyzed the collagen level in *Pkd-1* and *Pkd-2* conditionally deleted mouse kidneys. Strikingly, both *Pkd1* and *Pkd2* cKO kidneys, despite being highly cystic and enlarged, showed only spars amounts of collagen (blue staining) toward their end stage, by Masson’s trichrome staining (**Fig. S10A-B**). This was similar to *Ift88* cKO and *Tmem67* deficient kidneys but in great contrast to *Adamts9* cKO kidney collagen levels (**Fig. S9A**). Immunostaining for Collagen-IV showed a similar result, and moderately low interstitial staining were seen in *Pkd1* and *Pkd2* cKO kidneys (**Fig. S10C-D**). Quantitative RT-PCR analysis for *Col1a1* and *Col4a1* levels also corelated with these findings and showed either no change or downregulation of collagen transcript levels in *Pkd1* and *Pkd2* cKO kidneys (**Fig. S10E-F**). These findings greatly supported our theory that interstitial collagen level may be the deciphering factor governing kidney enlargement and type (nephronophthisis-like or ADPKD-like, **Fig. 10B**).

Since ADAMTS9 is not a collagenase, the increased collagen seen in *Adamts9* cKO kidneys is most likely due to loss of MT_1_-MMP cleavage (**Fig. 10A**). In our N-terminomics screen for identifying novel ADAMTS9 substrates, we also discovered that ADAMTS9 cleaves MT_1_-MMP [19]. MT_1_-MMP is a membrane anchored collagenase which is responsible for directly degrading type I, II and III, interstitial fibrillar collagens [52]. We previously showed that ADAMTS9-mediated cleavage releases the MT_1_-MMP catalytic domain from the cell surface, generating a soluble form of MT_1_-MMP [19]. Furthermore, MT_1_-MMP cell surface proteolytic activity triggers a cascade of events that can lead to deregulated collagen deposition (**Fig. 10A**). Cell surface MT_1_-MMP together with TIMP-2 (not shown) activates MMP2 by pro-domain cleavage and activated MMP-2 in tandem, activates MMP-9 [53, 54]. Both MMP2 and MMP9 are well-characterized collagenases [55, 56]. We previously showed that *ADAMTS9* KO cells and conditioned medium have increased collagenase activity due to increased cell surface activation of MMP2 and MMP9 due to loss of MT_1_-MMP cleavage [19]. Interestingly, MMP9 is an essential mediator of collagen deposition in the kidney, and mice lacking MMP-9 has reduced interstitial collagen levels in obstructive nephropathy [57]. This cascade of events leading to deregulated MMP activity triggered by loss of MT1-MMP cleavage (**Fig. 10A**), unique to *Adamts9* cKO kidneys but not the three other cystic kidney models, may be the distinguishing factor for the very high level of collagen seen in them (**Fig. S9A-E**). We propose here that this increased interstitial collagen may act as a very stiff “extracellular matrix scaffold”, mechanically preventing the *Adamts9* cKO cystic kidneys from enlarging, resulting in the distinctly small cystic kidney phenotype compared to the others (**Fig. 10B**).

The increased ECM may also act as an “extracellular reservoir” for signaling molecules and proinflammatory cytokines, explaining the comparatively high immunogenic response in the *Adamts9* cKO kidneys. The marked expansion of CD45⁺ leukocytes, CD11c⁺ dendritic cells, and F4/80⁺ macrophages that we observed in all models highlights a shared inflammatory signature across genetically distinct cystic kidney models. Immune cell recruitment has been well documented in polycystic kidney disease and nephronophthisis, where macrophages in particular have been implicated in promoting cyst growth, interstitial fibrosis, and tubular injury [51, 58]. Our findings extend these observations by demonstrating that loss of *Adamts9*, which has not previously been linked to renal immune dysregulation, is sufficient to generate a highly inflammatory cystic microenvironment, triggered by loss of cilia formation and interstitial ECM accumulation. Our results also suggest that cyst formation, regardless of the initiating genetic cause, may converge on common pathways, but utilizing different receptors for immune activation, which may be exacerbated by loss of extracellular matrix remodeling in the kidney. Defining the specific roles and activation states of the infiltrating immune populations, such as distinguishing pro-inflammatory versus reparative macrophage subsets, or assessing antigen-presenting functions of dendritic cells, will be crucial for designing cystic kidney sub-type specific immunotherapeutics. Testing whether modulation of inflammatory pathways can also modify cyst progression in *Adamts9* cKO kidneys in future studies may provide valuable insight into understanding the relationship of interstitial fibrosis and therapeutic outcome and determine whether the immune response is a driver of disease or a secondary consequence of cystic injury.

Our findings also highlight that, while high levels of active β-catenin leading to elevated canonical Wnt signaling may play a role in polycystic kidney disease etiology, it may not be the primary driver of cystogenesis in mouse kidneys. Since *Tmem67 11CLE* kidneys are highly cystic but have comparable levels of active β-catenin to that seen in a *Wt* mouse kidney. Prior to generation of the *Tmem67 11CLE* mouse model, it was unclear if the cystic kidney phenotype produced by *Tmem67*-null mice was due to loss of TMEM67 ciliary function or due to elevated Wnt signaling. We demonstrate here that highly cystic kidneys can be formed by the loss of TMEM67 function at the TZ, without upregulating active β-catenin levels. Although unlikely, it is unclear if elevated non-canonical Wnt signaling or ROR2 phosphorylation in *Tmem6711CLE* kidneys may somehow trigger a similar gene regulatory response, caused by increased canonical Wnt signaling seen in *Tmem67*-null kidneys, leading to cyst formation. Our findings strongly suggest that it is the loss of TMEM67 C-terminal half localization at the TZ, which is driving loss of ciliation or cilia function and triggers cyst formation in *Adamts9* cKO, *Tmem67*-null and *Tmem67 11CLE* kidneys. We have not further explored the signaling capabilities of the cleaved TMEM67 N-terminus which contains the CRD, in Wnt signaling augmentation in the mammalian kidney, since cystic kidneys are formed in its absence (*Tmem67 11CLE* kidneys) (**Fig. 10A**).

Since conditional deletion of versican and the versican-cleavage deficient mice have completely normal renal histopathology and lifespans, we conclude here that ADAMTS9-mediated versican proteolysis is unrelated to cystic kidney formation in mammals. However, it should be noted that, TMEM67 and MT_1_-MMP may not be the only targets of this crucial metalloproteinase in the kidney. Since loss of ADAMTS9 leads to a more severe loss of cilia phenotype (no cilia) while loss of TMEM67 or its cleavage leads to abnormal cilia formation (extremely long or short), it is most likely that ADAMTS9 may proteolytically cleave other proteins important for cilia formation in the kidney. A future proteomics study aimed at investigating the N-terminome of *Adamts9* cKO kidneys may shed valuable insight on uncovering these substrates.

Differential regulation of Laminin α5 in the four different cystic kidney models is also an interesting novel observation made by this study, and supports our theory on increased MT_1_-MMP activity in *Adamts9* cKO kidneys. Decreased Laminin α5 immunostaining in the GBM is unique to *Adamts9 cKO* kidneys, while the other three cystic kidneys significantly upregulated its expression and showed intense staining in the GBM. Which signaling pathways are responsible for this unique transcriptional upregulation of *Lama5* is unknown. Currently, there is no evidence to suggest Laminin α5 might be a direct substrate of ADAMTS9, which belongs to the chondroitin sulfate proteoglycanase subgroup of ADAMTS proteases. However, MT_1_-MMP has been previously shown to cleave the alpha-5 chain of purified Laminin-10 [59]. We therefore suggest increased cell surface MT_1_-MMP present in *Adamts9* cKO kidney podocytes leads to increased degradation of Laminin α5 in the GBM resulting in its decreased staining in *Adamts9* cKO kidneys, while transcriptional upregulation of *Lama5* leads to increased Laminin α5 levels in *Ift88* cKO and *Tmem67* deficient kidneys. It is unclear if increased MT_1_-MMP mediated catabolism of Laminin α5 contributes to the *Adamts9* cKO renal pathology since genetic deletion of *Lama5* leads to cystic kidney formation in mice without affecting ciliogenesis [28, 29].

While we utilized *Six2*-Cre mediated conditional deletion of *Ift88* as a control model lacking primary cilia in this study, it is also important to note that this is the first study to report the outcome of deleting *Ift88* in the nephrogenic lineage specifically. Intriguingly, loss of *Ift88* in the ureteric lineage, proximal tubules or postnatally by inducible Cre expression throughout the kidney, leads to relatively slow onset cystic kidney formation [26, 60, 61]. Our results suggest that nephrogenic lineage deletion is comparably more severe and results in fast onset cystic kidney development and lethality of *Ift88* cKO mice by 14 days. Furthermore, E18.5 *Ift88* cKO kidneys although show onset of tubular enlargements, had otherwise normal kidney histopathology with no change in glomerular numbers or nephric differentiation, strengthening the notion that the cilium is indeed dispensable for early events of kidney development in mammals.

The factors which regulate differential *Adamts9* and *Tmem67* expression in male versus female proximal tubules are unknown. Whether simply higher levels of *Adamts9* expression in male proximal tubules causes male *Adamts9* cKO kidneys to have a more severe pathology, when deleted, is unknown. Since both *Tmem67*-null and *11CLE* mice do not show a sex bias, the male sex specific phenotypic burden is unique to *Adamts9* loss of function. Since *Six2*-Cre mediated deletion of *Ift88* does not show a sex bias, and kidneys from both sexes were equally affected, expression of *Six2*-Cre is most likely not sex specific in the murine kidney. It is possible that *Adamts9* expression may be differentially regulated in male and female cKO kidneys, downstream of the kidney injury response, which strikingly has gender specific gene expression differences in both humans and mice [62]. It will be interesting to investigate if *ADAMTS9* ciliopathy patients show any sex specific bias in survival and renal pathology and function in a future clinical study. Based on the small cohort of *ADAMTS9* patients identified to date and the unavailability of extensive renal histopathological data, it is unclear if *ADAMTS9* NPHP patients also have a sex specific cystic kidney severity.

Finally, when the very first patients carrying *ADAMTS9* mutations were discovered [14], the fact that they were ciliopathy patients, diagnosed with NPHP and JBTS was a puzzling new development. This was because ADAMTS9 was a *bona fide* extracellular matrix metalloproteinase, which has been extensively studied for decades prior, in relation to ECM homeostasis, and had no apparent connection to the cilium or ciliary function, known to exclusively act outside of the cell. The concurrent development of *Adamts9* floxed mice and hypomorphic mice (*Adamts9^Gt^)* which allowed us to study the function of ADAMTS9 beyond gastrulation in mice, and identification of novel substrates utilizing N-terminomics have now solidified ADAMTS9 as a crucial extracellular protease for ciliogenesis. Here, building upon these recent discoveries, we show that indubitably, loss of this crucial enzyme leads to NPHP in mammals and uncover the unique underlying molecular mechanism.

## ACKNOWLEDGMENTS

This work was funded by the National Institute of Health grant 5R01DK126804 and The Worcester Foundation grant awards to SN. This work utilized SEM, TEM and Ultramicrotomy equipment purchased from NIH awards S10RR021043, S10OD025113, and SI0OD021580. A TissueGnostics-SL slide scanner used in the study was funded by a Massachusetts Life Science Center, Bits to Bytes grant. *Pkd1^fl/fl^;HoxB7-Cre* and *Pkd2^fl/fl^*;*Rosa26-Cr*e^ERT^ paraffin-embedded kidney blocks and frozen kidneys were obtained from the Kansas PKD Research and Translational Core Center at KUMC (U54 DK126126) and the PKD Research Resource Consortium (PKD RRC). We thank professors George Witman and Greg Pazour for their valuable input and thank Dr. Suneel Apte, Chuck Frevert and Tom Wight for developing *Adamts9* and *Vcan* mouse lines and Dr. Colin Johnson for the TMEM67-C antibody utilized in this study.

## AUTHOR CONTRIBUTIONS

S.F., M.A., K.R., G.I.K., P.U.A. and S.N. conceived and designed the experiments. S.F., M.A., K.R., G.I.K., M.A.K, W.W. and S.N. performed experiments. S.F., M.A., K.R., G.I.K., P.V.T., P.U.A., and S.N. wrote the manuscript. All authors read, edited and approved the manuscript.

## COMPETING INTERESTS

The authors declare no competing interests

## METHODS

### Contact for reagent and resource sharing

Requests for resources, reagents, and further information should be directed to and will be fulfilled by the lead Contact, Dr. Sumeda Nandadasa

### Data availability

All data that support the findings of this study are available from the corresponding author upon reasonable request.

### Mice

All animal studies were performed with approval of the institutional animal care and use committees (IACUC) of the UMass Chan School of Medicine and University of Kansas Medical Center. The mice were housed under a 12-h light and 12-h dark cycle and had unrestricted access to food and water. The *Tmem67^+/-^* mice (C57BL/6NJ-*Tmem67em1^(IMPC)J/Mmjax^*) were purchased from the Jackson Laboratory (JAX, 051248). *Tmem67^ΔCLE/+^* founder mice were generated in the C57BL/6J background by Cyagen (Santa Clara, CA) as previously described [20]. The *Ift88^Fl^* mice were purchased from Jackson Laboratory (JAX: 022409). *Adamts9^Fl/+^* and *Adamts9^Del/+^* mice [22, 63] were obtained from Dr. Suneel Apte and were maintained in the C57BL/6J background. *Six2*-Cre eGFP mice were obtained from the UT Southwestern, George. M O’Brien Kidney Research Core Center, and were maintained in a mixed background of C57BL/6J, Swiss Webster and FVB [23]. *Ksp*-Cre mice were purchased from Jackson Laboratory (JAX: 012237). *Vcan^Fl/+^* mice (*Vcan^tm1.1Cwf^*) [64], were provided by Dr. Charles Frevert and Thomas Wight. *Vcan^Hdf/+^* (null-allele) and *Vcan^AA/+^*mice [27, 65] were provided by Dr. Suneel Apte and were maintained in the C57BL/6J background. *Pkd1^Fl^*, *Pkd2^Fl^*, *HoxB7-Cre* and *ROSA26-Cre^ERT^*mice were purchased from Jackson Laboratory (JAX, 010671, 017292, 004692, 004847) and maintained on a C57BL/6J background. *Pkd1* cKO mice were analyzed at P14. To generate *Pkd2* cKO; *ROSA26-CreERT* mice, nursing mothers were injected intraperitoneally with tamoxifen (8mg/40g; Sigma-Aldrich, T5648) at P0 to induce gene deletion. Offspring were analyzed at P21. A list of genotyping primers used to genotype each mouse line is provided in **Table S1**.

### Cell Culture

Primary kidney cell cultures were established from mouse kidneys at postnatal day 14 or 21. In brief, kidneys were finely minced and digested in 1.5% Collagenase IV for 1 hour (hr) at 37°C. Cells were cultured in media containing DMEM/F12 (Gibco;11330-032;), 2% FBS, 200 U/mL penicillin/streptomycin (Gibco; 15140-122), glutamine, 5 mg/L human transferrin, 5 mg/L recombinant human insulin, 7.5 nM sodium selenite, and 2nM 3,3,5’-Triiodo-Thyronine 2 nM at 37°C, 5% CO2 as previously described [66]. Wild type HTERT-RPE-1 (ATCC, CRL-4000) and *ADAMTS9* KO RPE-1 cells [15], were maintained in DMEM/F12 (Gibco;11330-032;) with 10% FBS and 200 U/mL penicillin/streptomycin (Gibco; 15140-122) and maintained at 37°C with 5% CO_2_.

### Plasmid DNA constructs and transfections

The ADAMTS9 S636F, R647Q, R1203W, C1024*, P1665S and S1826N patient variants were generated using the Q5 Site-Directed Mutagenesis Kit (NEB; E0554S) using the full-length ADAMTS9 plasmid construct in the pCDNA3.1Myc/His backbone described previously [15] utilizing the primers listed in **Table S2**. T65R and Q1525Hf/s variants were previously generated in the pENTER backbone [14]. All constructs were verified through Sanger sequencing. For plasmid DNA transfections, RPE-1 cells were transfected with 300 ng of plasmid DNA per well in 8-chamber slides (Corning; 354118) grown at 60-80% confluency, using PEI MAX transfection reagent (Polysciences; 24765). Transfected cells were washed by PBS and serum starved for 24 hrs by culturing in serum-free DMEM/F12 culture medium, for ciliogenesis and TZ analysis.

### Western blot analysis

Confluent primary kidney cell cultures or RPE-1 cultures in 6-well plates were serum starved for 48 hours in 1 ml of serum-free DMEM/F12 culture medium and lysed using 300 μL Transmembrane lysis buffer (FIVEphoton biochemicals; TmPER-50) as previously described [20]. 6x Laemmli sample buffer was used for boiling cell lysates or conditioned medium for 10 min at 100 °C. Samples were loaded on to 7.5% acrylamide gels and electrophoresed for 60-90 minutes at 110V. Protein bands were transferred to Immobilon-FL transfer membranes (EMD Millipore, IPFL00010) for 3 hrs at 333 mA at 4 °C, and subsequently blocked with 5% nonfat dry milk dissolved in PBSTw (PBS+ 0.1% Tween-20) for 1 hour. Primary antibodies were diluted in 5% nonfat milk and incubated overnight at 4 °C with agitation. Li-Cor fluorescent secondary antibodies were diluted in PBSTw and incubated for 2 hrs at room temperature (RT) following 3x PBSTw washes. Western blot membranes were imaged using the LI-COR Odyssey M scanner. Western blot band intensities were quantified using NIH Image J (FIJI). A list of all antibodies and dilutions used in western blotting are provided in the key resource table (**Table S4)**.

### Immunostaining and fluorescent microscopy

Dissected mouse kidneys were bisected and fixed overnight at 4°C in 4% PFA. For cryo sectioning, immediately following 3x PBSTw washes, fixed samples were placed in 15% sucrose overnight followed by 30% sucrose at 4°C overnight. Samples were imbedded in OCT (Tissue-Tek; 4583). Cryo-embedded kidneys were sectioned via a Leica CM 1950 cryostat collecting 7 μm frozen sections. For paraffin sectioning, fixed kidneys were washed 3x in PBSTw following fixation and paraffin embedded using standard procedures. A Leica RM2155 motorized microtome was used for collecting 7 μm paraffin sections. Deparaffinized paraffin sections and rehydrated cryo sections in PBS were blocked with 5% NGS in PBSTx (PBS+ 0.1% Triton X-100) for 2 hrs. Kidney sections were incubated overnight at 4°C with primary antibodies diluted in blocking buffer. Sections were washed 3x with PBSTx and incubated in secondary antibodies diluted in PBS for 2 hrs in the dark at RT. Following 3x PBSTx washes, the slides were mounted in Prolong gold antifade mounting media containing DAPI (Invitrogen; P36931). For immunostaining primary kidney culture and RPE-1 cells, serum starved cells grown in 8-chamber slides (Corning; 354118) were fixed with ice cold methanol for 5 minutes at -20°C for TZ analysis or 4% PFA for 30 minutes at RT for cilium staining. Cells were washed 3x with PBSTw and blocked for 2 hrs in 5% NGS made in PBSTw. Primary antibodies were diluted in blocking buffer and incubated overnight at 4°C with agitation. After 3x PBSTw washes, the cells were incubated with secondary antibodies diluted in PBSTw for 2 hrs at RT. After 3x PBSTw washes, the slides were mounted with Prolong gold antifade mounting medium containing DAPI (Invitrogen, P36931), and curated overnight at RT. Confocal images were obtained using a Leica SP8 laser scanning confocal microscope with a 63x 1.47 N.A. oil-immersion objective, a 40x N.A. 1.30 oil objective, or a 20x N.A. 0.75 dry objective. Whole kidney fluorescent images were obtained utilizing a TissueGnostics TIssueFAXS microscope. A list of all antibodies and dilutions used in immunostaining are provided in the key resource table (**Table S4)**.

### Cystic kidney histopathology assays

Dissected mouse kidneys were bisected using a sharp razor blade and fixed overnight at 4°C in 4% PFA and then washed 3x with PBSTw. Fixed kidney samples were then dehydrated over two consecutive nights, in 50% ethanol and then 70% ethanol at 4°C. Kidney samples were embedded in paraffin using a standard tissue processing procedure. Kidney sections were cut at 7 μm using a Leica RM215 motorized microtome. For hematoxylin and eosin staining, sections were deparaffined and stained with eosin Y 1% (VWR; 10143-130) and hematoxylin (VWR; 10143-142) and mounted in Cytoseal XYL medium (Adwin Scientific, NC9527349). For Masson’s Trichrome staining, kidney sections were deparaffined and refixed in Bouin’s fixative (Electron Microscopy Sciences (EMS); 26367-01), stained with Weigert’s iron hematoxylin A & B solutions (EMS; 26367-02 and -03), Biebrich scarlet solution (EMS; 26367-04), and aniline blue solution (EMS; 26367-06) followed by a 5 minute 1% acetic acid (EMS; 26367-07) fixation. For Periodic Acid Schiff (PAS) staining, slides were deparaffined followed by staining with 1% periodic acid (A223-25, Fisher), Schiff’s reagent (26853-01, Electron Microscopy), and Mayer’s hematoxylin (Volu-Sol, VMH032). All samples were mounted in Cytoseal XYL medium and imaged using a TissueGnostics TIssueFAXS slide scanner or a Zeiss Axioplan microscope equipped with a Leica DM6200 camera.

### Quantitative real time PCR (qRT-PCR)

mRNA was harvested from whole mouse kidneys snap frozen in 500 μL Trizol reagent and crushed into a fine powder using a frozen pestle and motor. The thawed lysates was sonicated for 30 seconds followed by chloroform extraction and isopropanol precipitation. RNA pellets were washed with 100% and 70% ethanol and resuspended in nuclease-free water. 2 μg of mRNA per sample was used in cDNA synthesis using the High Capacity cDNA Reverse Transcription Kit (ThermoFisher; 4368814). A BioRad CFX connect real time PCR machine was used in combination with Bullseye EvaGreen qPCR master mix (Midsci; BEQPCR-R) for determining ΔCT values. Each sample was run in duplicate with at least 4 experimental replicates per group. All gene expression data was normalized to *18s*. A list of qRT-PCR primers used in this study are provided in **Table S3.**

### Transmission and Scanning Electron Microscopy

All EM samples in this study were processed and analyzed at the University of Massachusetts Chan Medical School Electron Microscopy core facility according to standard procedures. In brief, mouse kidneys were fixed in 2.5% glutaraldehyde with 1.6% paraformaldehyde in 0.1 M sodium cacodylate buffer (pH 7.2), overnight at 4 °C. The samples were then washed 2x in fresh fixation buffer at RT and post-fixed with 1% osmium tetroxide for 1 hr at RT. Samples were then washed 2x with dH_2_O for 5 min and dehydrated using a graded ethanol series (of 20% increments) and finally in two changes of 100% ethanol. Fully dehydrated samples were infiltrated with 100% propylene oxide and then with propylene oxide followed by a SPI-Pon 812 50%–50% mixture overnight. Samples were polymerized at 68 °C in plastic capsules immediately following 3x fresh 100% SPI-Pon 812 resin washes. Polymerized samples were oriented, and 70 nm ultra-thin sections were collected using a Leica EM UC7 ultramicrotome equipped with a Diatome ultra 45 diamond knife for TEM analysis. Sections were placed on copper support grids and contrasted with lead citrate and uranyl acetate. TEM analysis was conducted utilizing a FEI Tecnai 12 BT with 120 kV accelerating voltage, and images were obtained by a Gatan TEM CCD camera. For freeze-fracture SEM analysis of wild type and cystic kidney cilia, fixed kidneys were dehydrated through a graded series of ethanol and either directly critical point dried or immersed in liquid nitrogen and placed on a liquid nitrogen-cooled block, and fractured using a pre-cooled hammer and razor blade. Fractured kidney pieces were collected and thawed in 100% ethanol and critical point dried. Kidney samples were mounted on aluminum SEM stubs with fracture surfaces pointing up and metal-coated (Au/Pb, 60/40) prior to imaging using a FEI Quanta 200 MKII FeSEM.

### Flow Cytometry

Immediately following euthanasia, freshly harvested kidneys were chopped with a razor blade and placed into collagenase-containing DMEM with 10% FBS for 1 hr at 37C. The samples were then mashed through 40 micron mesh filters (Fisher Scientific; 22363547) and passed through a 27G syringe prior to centrifugation at 1,500g for 5 min. Samples were resuspended in 1 mL ACK lysis buffer (Thermo Fisher; A1049201) for 10 min followed by an additional wash with 1 mL ACK lysis buffer. The concentration of cells were determined using a Countess 3 cell counter (Invitrogen) and resuspended in FACS buffer, and plated at 1 million cells per well in 96-well clear round-bottom plates for immunostaining. Fc block (BD Biosciences; 553142) diluted 1:50 in FACS buffer was added prior to staining at 4C for 30 min using 1 μg per well of each of the following fluorescently tagged primary antibodies in 100 μl FACS buffer. All primary antibodies and their dilutions used in flow cytometry assays are provided in the key resource table (**Table S4**). Cells were stained with DAPI (1:1000 in FACS buffer), then washed 3x with PBS and resuspended in PBS before analysis on a ZE5 Cell Analyzer (Bio-Rad Laboratories). Flow cytometry data was analyzed using FlowJo software.

### Quantification of renal histopathology and immunostaining data

Kidney area, glomeruli counts, cilia frequency, cilia length, transition zone intensity, active β-catenin intensity were all quantified using NIH Image J (FIJI). The cell counter function was used to quantify the number of glomeruli per midline kidney section and cilia frequencies. The line trace function was used to measure cilia lengths. The kidney area was quantified using the polygon tool. For quantifying transition zone marker intensities, the mean pixel intensity was measured from a ROI, specifying the signal directly above the γ-tubulin staining as previously described [20]. For active β-catenin staining, the polygon tool was used to trace the kidney tubule and measure the mean pixel intensity in the active β-catenin signal (568/red). All statistical tests were done using GraphPad Prism software with statistical tests and significance indicated in the figure legends.

## SUPPLEMENTAL FIGURE LEGENDS AND MOVIE CAPTIONS.

**Figure S1.**
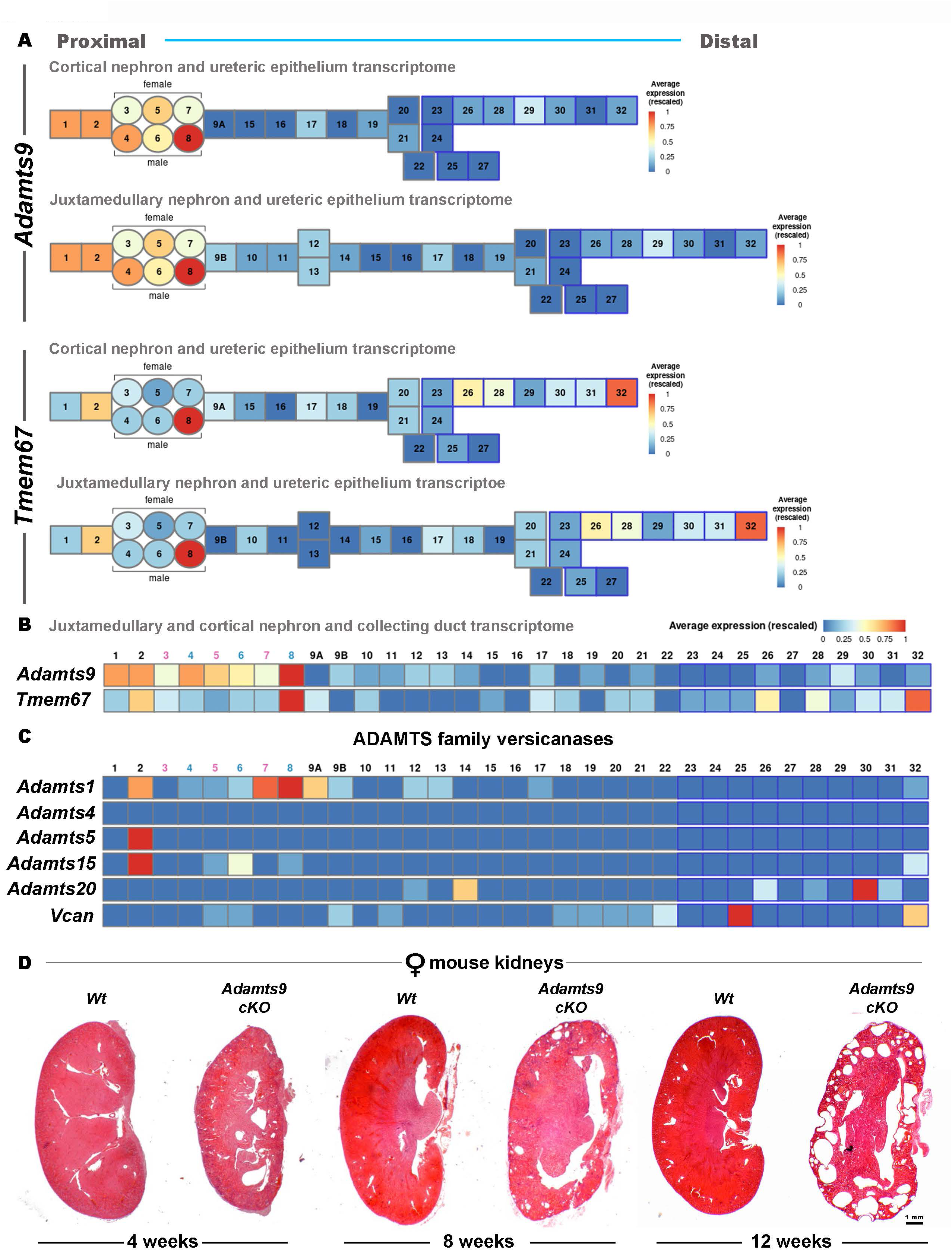
Sex specific expression patterns of *Adamts9* and *Tmem67* in the murine kidney and delayed onset of cystic kidneys in *Adamts9* cKO females. (**A-B**) Single cell RNA sequencing data for *Adamts9* and *Tmem67* in the murine kidney comparing male and female adult mouse kidneys utilizing The Kidney Cell Explorer (https://cello.shinyapps.io/kidneycellexplorer/). Ontology of cell populations pertaining to *Adamts9* and *Tmem67* expression: 1-podocytes, 2- Parietal epithelium, 3- Segment 1 of proximal tubules (female), 4- Segment 1 of proximal tubules (male), 5- Segment 2 of proximal tubules (female), 6- Segment 2 of proximal tubules (male), 7- Segment 3 of proximal tubules (female), 8- Segment 3 of proximal tubules (male), 26- Principal cell of outer medullary collecting duct, 32- Deep medullary epithelium of pelvis. (C) Expression pattern of other versicanases belonging to the ADAMTS family and *Vcan* in male and female adult mouse kidneys. Ontology of key cell populations: 2- Parietal epithelium, 3- Segment 1 of proximal tubules (female), 4- Segment 1 of proximal tubules (male), 5-Segment 2 of proximal tubules (female), 6- Segment 2 of proximal tubules (male), 7- Segment 3 of proximal tubules (female), 8- Segment 3 of proximal tubules (male), 9A- Loop of Henley, thin descending limb of inner stripe of outer medulla of cortical nephron, 14- Upper loop of Henley, thin ascending limb of inner medulla of juxtamedullary nephron, 25- Intercalated type A cell of outer medullary collecting duct, 32- Deep medullary epithelium of pelvis. (D) Hematoxylin and eosin staining of *Adamts9* cKO and *Wt* female littermate kidneys showing relatively slow onset of cystic pathology during 4-12 weeks of age.

**Figure S2.**
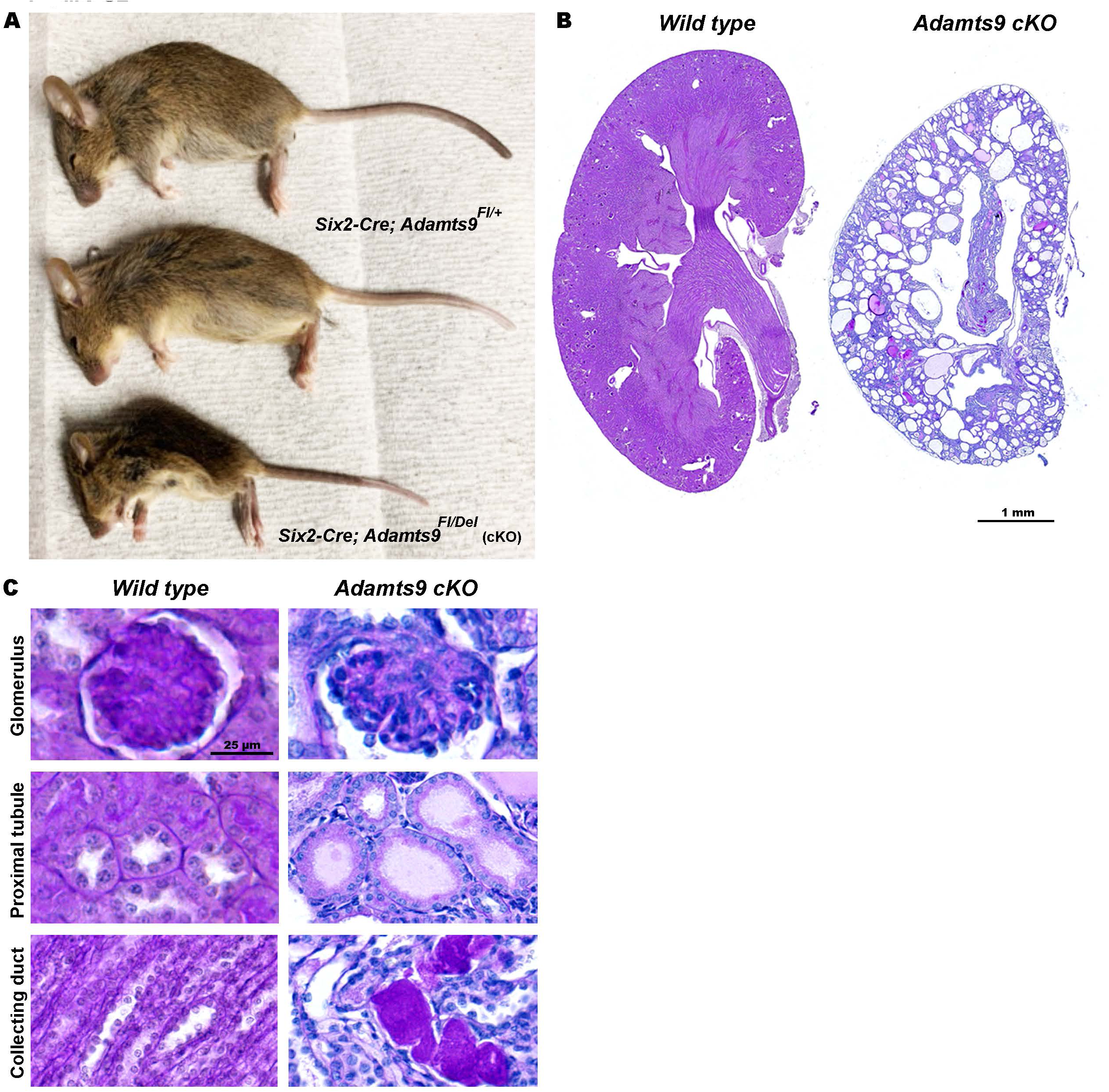
End stage male *Adamts9* cKO mice and Periodic Acid-Schiff (PAS) staining of kidneys. (**A**) *Adamts9* cKO male mice become distinguishably smaller from heterozygous littermates at 4 weeks of age. (**B-C**) PAS staining of p21 *Wt* and *Adamts9* cKO male kidneys with magnified images of glomeruli, proximal tubules, and collecting ducts (n=3 kidneys from each genotype). Scale bar in **B** is 1mm and 25 µm in **C**.

**Figure S3.**
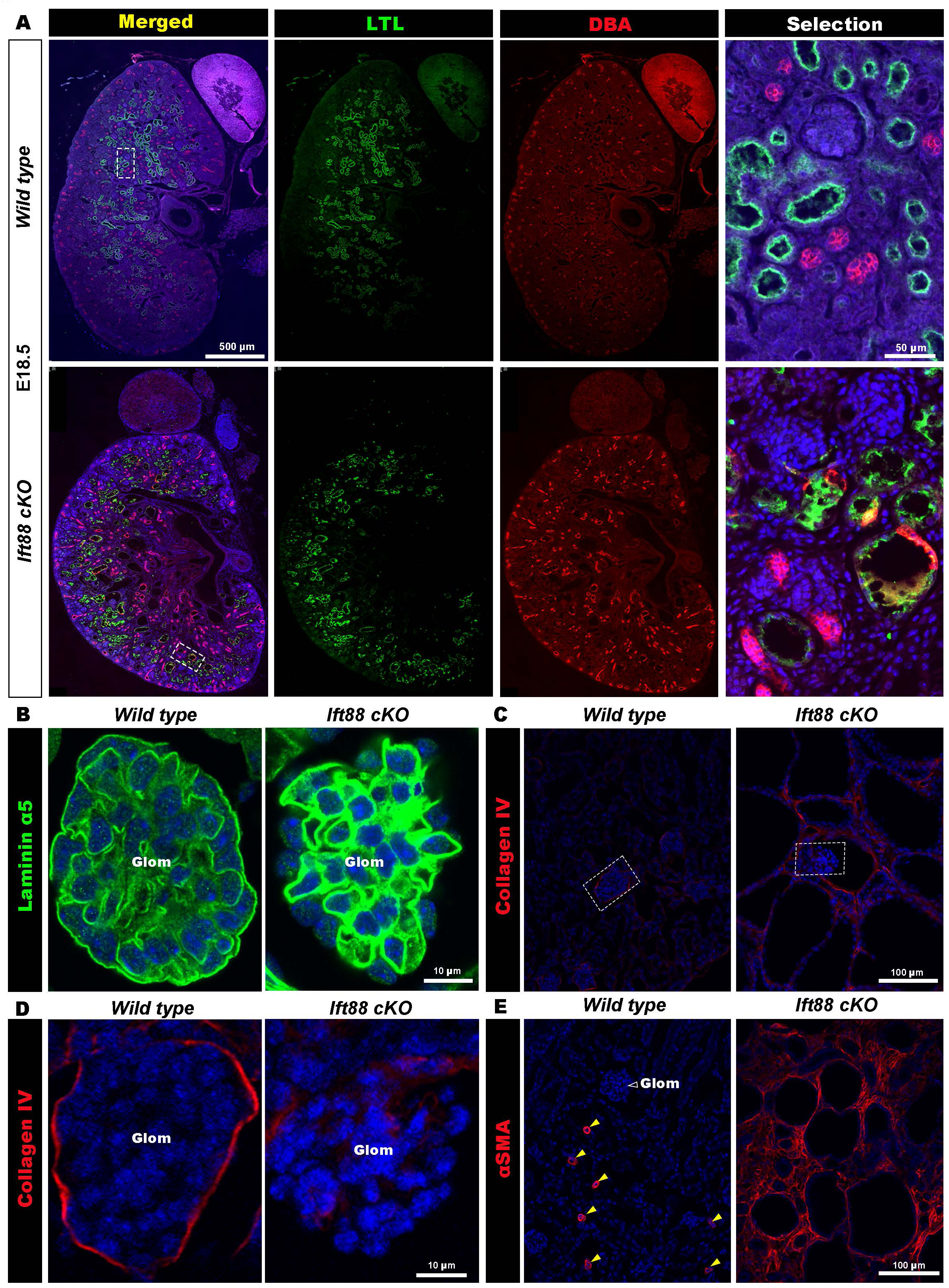
Normal embryonic nephrogenesis and ECM homeostasis defects in *Ift88* cKO kidneys. (A) Male E18.5 kidneys immunostained for proximal tubules (LTL, green) and collecting ducts (DBA, red) show comparable staining levels in *Wt* and *Ift88* cKO kidneys. The boxed medullary region shown in high magnification show enlarged proximal tubule morphologies in *Ift88* cKO kidneys. (B) Intense laminin α5 (green) immunostaining in *Ift88* cKO GBM in comparison to *Wt* p14 kidneys (n=3 male kidneys/ genotype). (**C-D**) Collagen IV (red) immunostaining which is limited to the glomerulus in *Wt* kidneys did not show highly elevated staining in the *Ift88* cKO kidneys. Glomeruli marked by the white box in **B** are shown in high magnification in **C** (n=3 male kidneys/ genotype). (E) α-SMA (red) immunostaining which is limited to the vasculature in *Wt* kidneys (yellow arrowheads) showed intense staining throughout the *Ift88* cKO kidneys. White open arrowhead indicates a glomerulus (n=3 male kidneys/ genotype).

**Figure S4.**
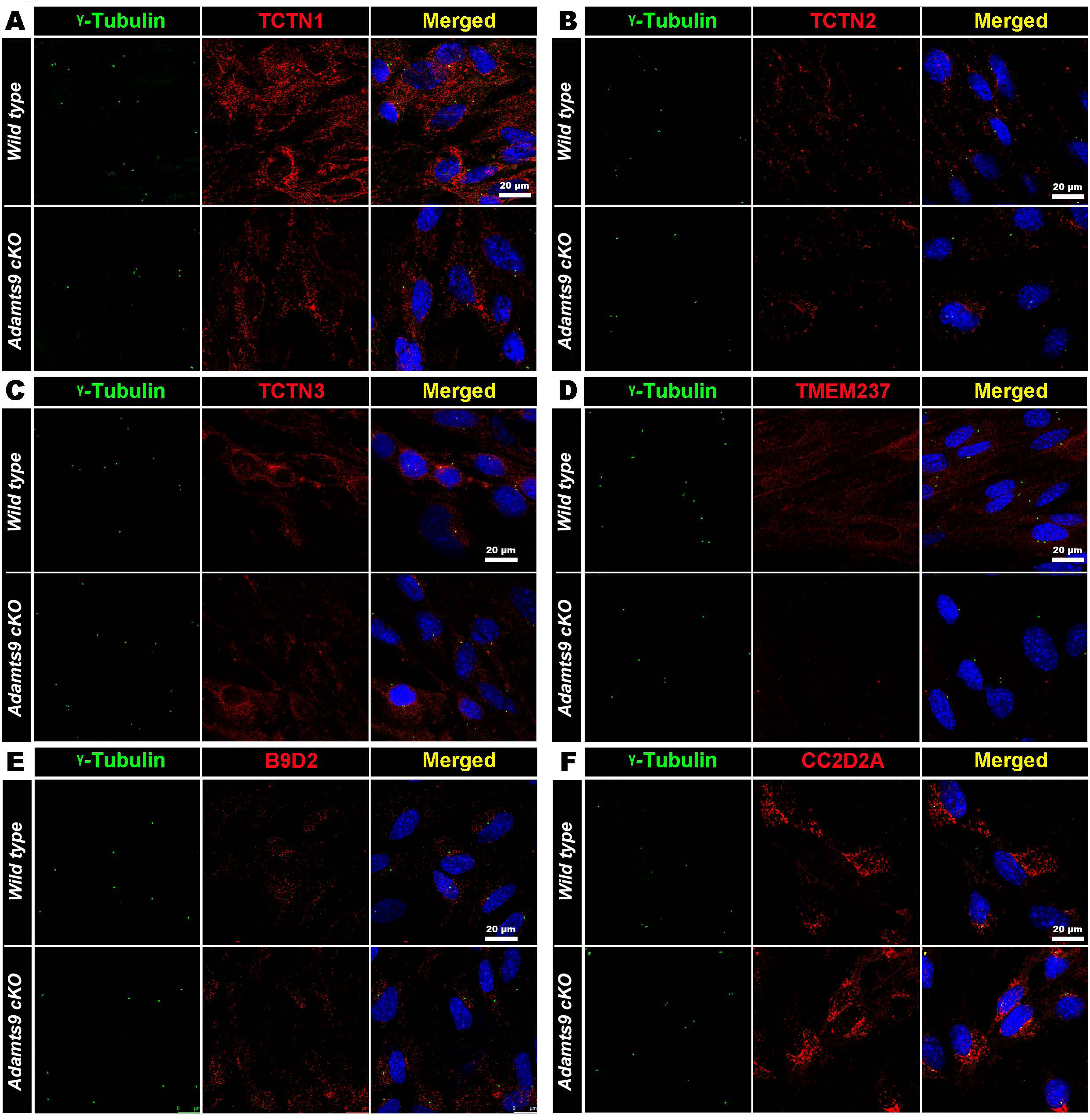
Low magnification images of transition zone MKS/B9 module immunostaining of ice-cold methanol fixed primary kidney cultures of *Wt* and *Adamts9* cKO kidneys. *Adamts9* cKO kidney primary cell cultures immunostained for the basal body (γ-tubulin, green) and the indicated transition zone markers (red, **A-F**). Scale bars are 20 µm in **A-F**.

**Figure S5.**
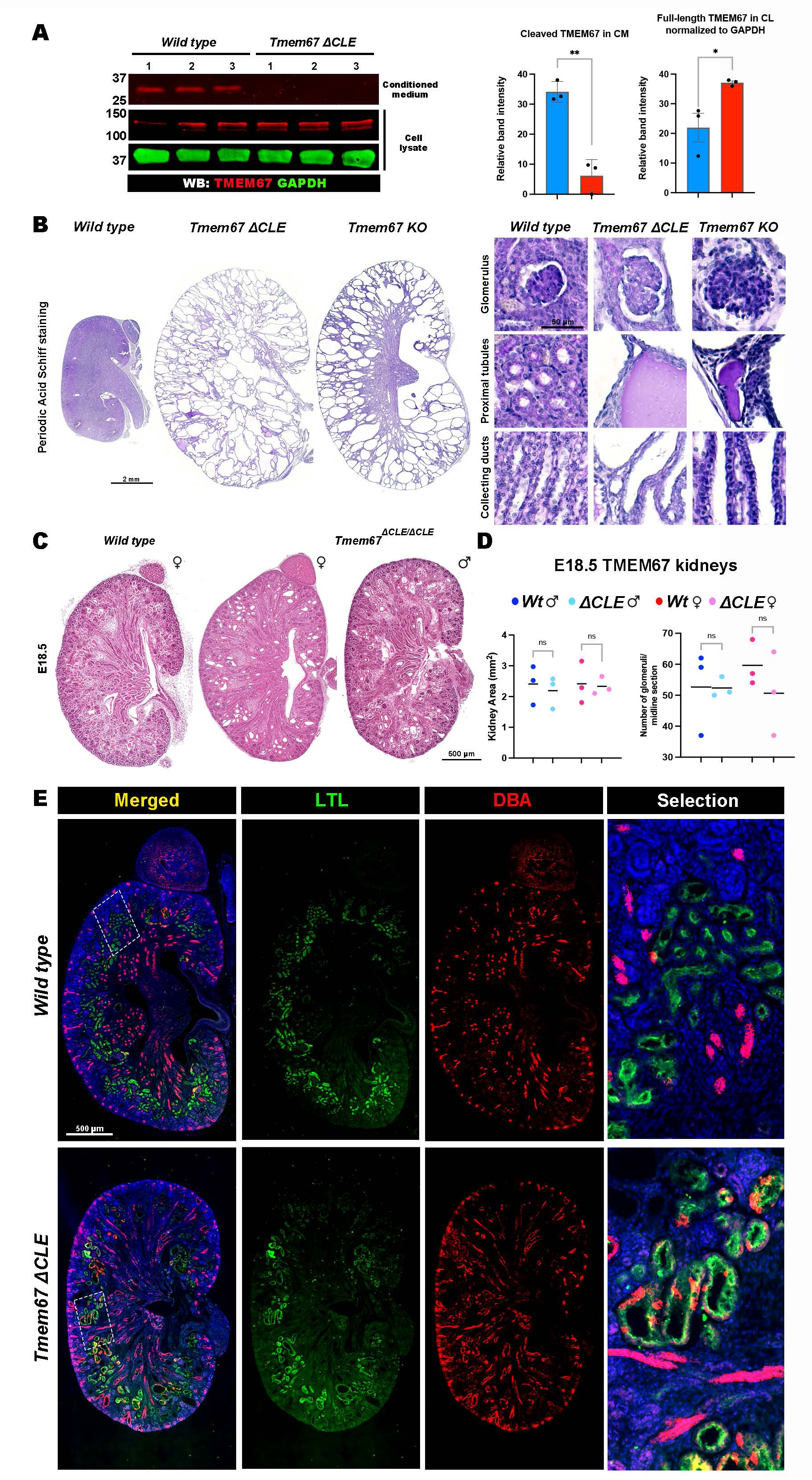
TMEM67 extracellular domain cleavage deficient mice (*Tmem67^ΔCLE^/ ^ΔCLE^*) undergo normal embryonic kidney development. (A) Western blot analysis of TMEM67 cleavage in *Wt* and *Tmem67^ΔCLE/ΔCLE^* primary kidney cultures show loss of the 34 kDa N-terminal fragment generated by ADAMTS9-mediated TMEM67 extracellular domain cleavage in the conditioned medium and increased full-length TMEM67 protein in the cell lysates of *Tmem67^ΔCLE/ΔCLE^*kidneys. (n=3 independent primary cultures, TMEM67 (red) and GAPDH (green). Error bars indicate Mean ± SEM, ** indicates a *p*-value <0.01 and * indicates a *p*-value <0.05 calculated by two-tailed unpaired *t*-test. (B) PAS staining of p14 *Wt, Tmem67 KO* and *Tmem67^ΔCLE/ΔCLE^* with magnified images of the glomeruli, proximal tubules, and collecting ducts (n=3 male kidneys/ genotype). (C) Hematoxylin and eosin staining of E18.5 kidneys show relatively normal histology and onset of cyst formation in both male and female *Tmem67^ΔCLE/ΔCLE^* kidneys (n=3 kidneys/ sex/ genotype). (D) Quantification of kidney area and number of glomeruli show normal nephrogenesis of E18.5 *Tmem67^ΔCLE/ΔCLE^*kidneys. (n=3 kidneys/ sex/ genotype). Lines indicate Mean, ns indicates a p-value >0.05 calculated by two-tailed unpaired *t*-test. (E) E18.5 kidneys immunostained for proximal tubules (LTL, green) and collecting ducts (DBA, red) show comparable staining levels in *Wt* and *Tmem67^ΔCLE/ΔCLE^* kidneys. The boxed medullary region shown in high magnification show enlarged proximal tubule morphologies in *Tmem67^ΔCLE/ΔCLE^* kidneys. Scale bars are 2 mm and 50 µm in **B**, 500 µm in **C** and **D**.

**Figure S6.**
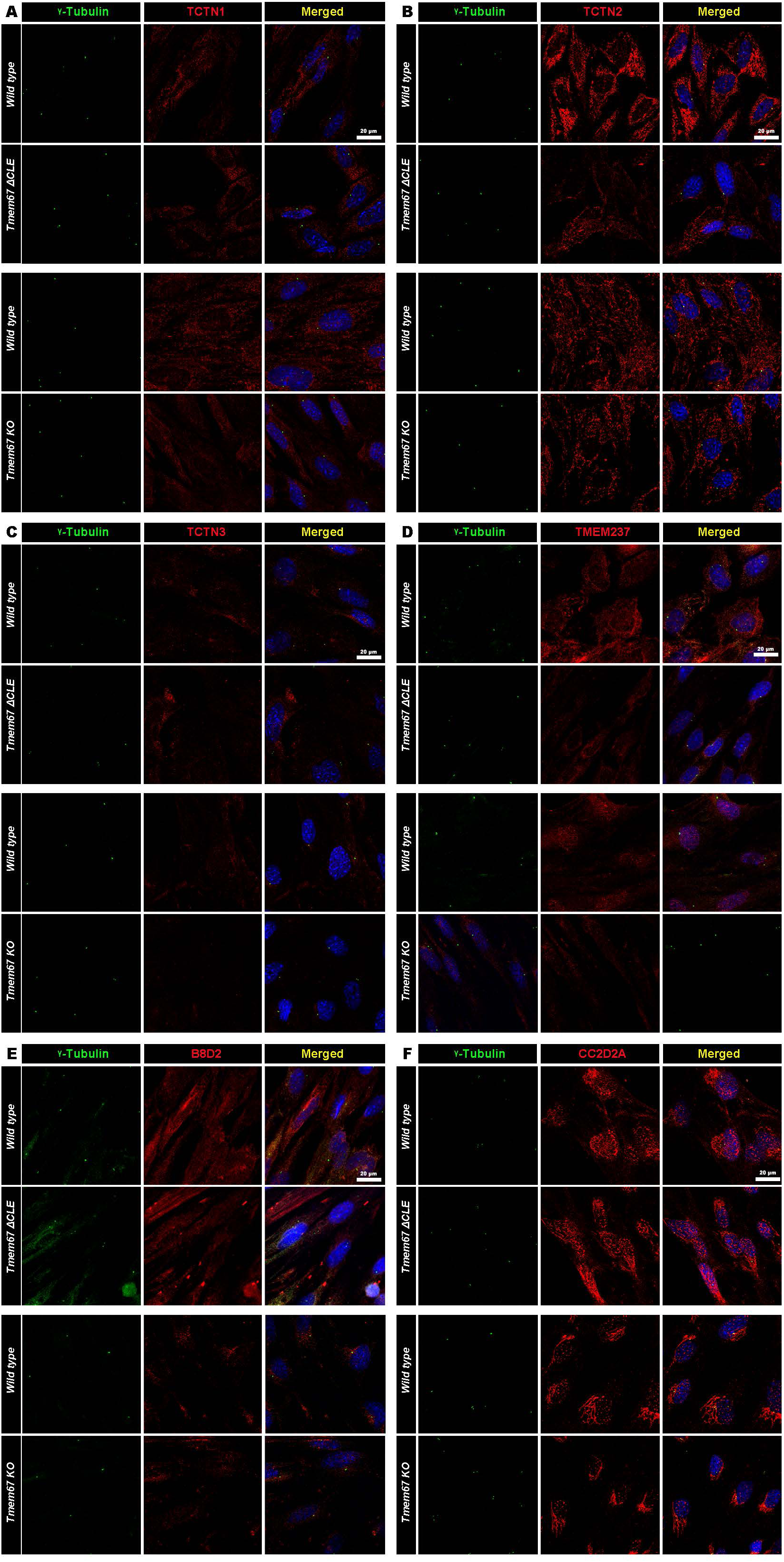
Low magnification images of transition zone MKS/B9 module immunostaining of ice-cold methanol fixed primary kidney cultures of *Wt*, *Tmem67 KO* and *Tmem67 ΔCLE* kidneys. *Wt*, *Tmem67 KO* and *Tmem67 ΔCLE* kidney primary cell cultures immunostained for the basal body (γ-tubulin, green) and the indicated transition zone markers (red, **A-F**). Scale bars are 20 µm in **A-F**.

**Figure S7.**
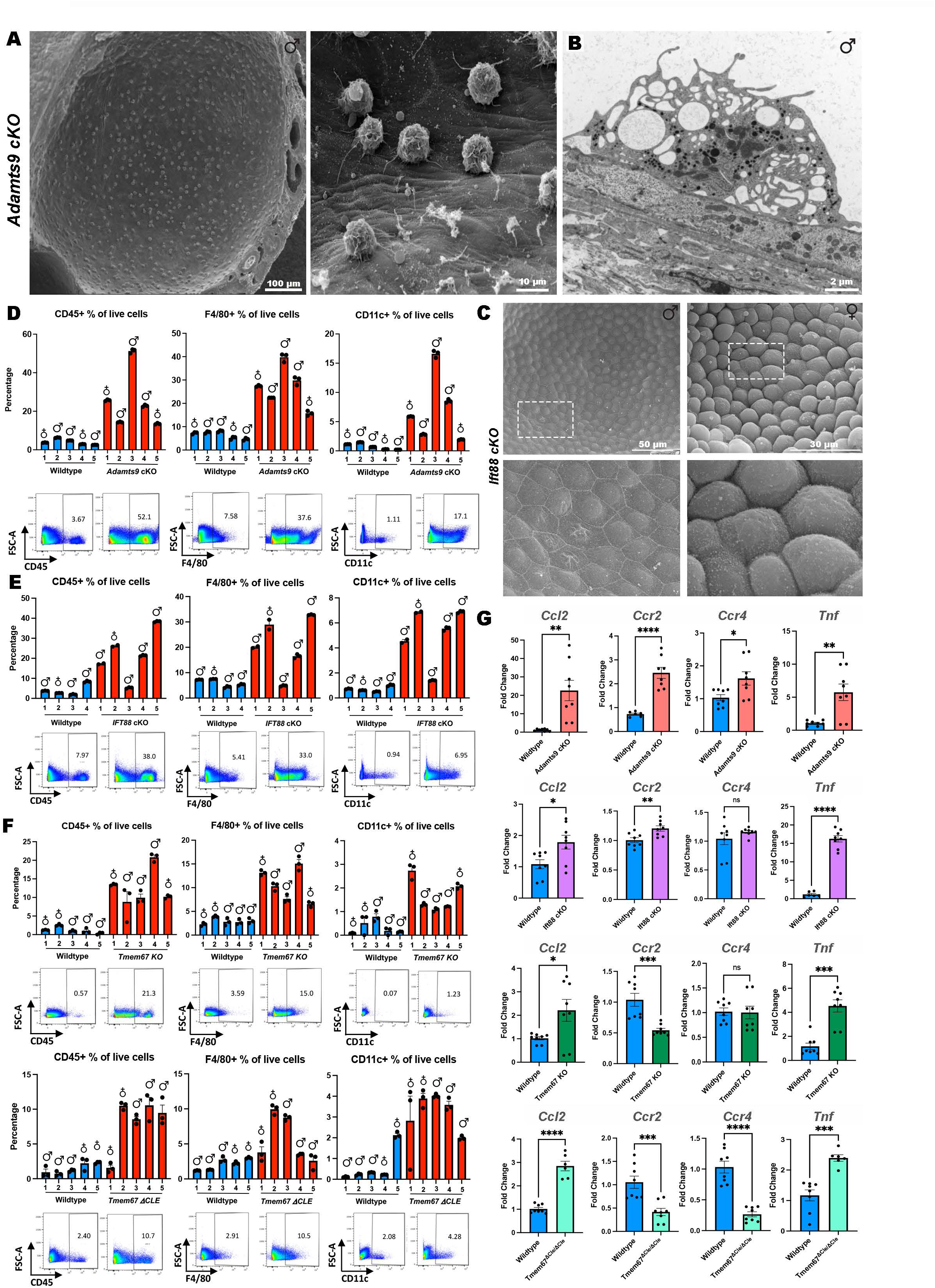
Macrophage infiltration in *Adamts9* cKO kidney cystic lumens and immune response of cystic kidney models. (**A-B**) SEM (**A**), and TEM (**B**) images of p21 male *Adamts9* cKO kidneys showing large numbers of macrophages residing in their cystic lumens (n=3 kidneys). Left image in **A** shows a single cystic lumen from freeze-fracture SEM and the right image showed a high magnification of the cystic surface. TEM image in **B** shows morphology of the macrophages residing in the cystic lumen with many membrane protrusions, vacuoles and lysosomes. (C) Freeze-fracture SEM images of p14 male and female *Ift88* cKO kidney cysts which did not show any macrophages residing in their cystic lumens (n=3 kidneys). (**D-F**) Flow cytometry analysis of immune cells present in whole kidneys of *Adamts9* cKO (**D**), *Ift88* cKO (**E**), *Tmem67* KO and *Tmem67^ΔCLE/ΔCLE^* kidneys (**F**), in comparison to littermate kidneys show high levels of CD45+, F4/80+ and CD11c+ cells in cystic kidneys from both sexes with the *Adamts9* cKO kidneys showing the highest percentages for each group (n=39 mice/ kidneys). Blue boxes mark *Wt* littermates and red boxes mark cystic kidneys. Error bars in each column indicate Mean ± SEM of three technical replicates of each kidney. Scatter plots of a representee male kidney is shown for each genetic group. (**G**) qRT-PCR analysis for expression levels of *Ccl2, Ccr2, Ccr4 and Tnf* in male cystic kidneys of *Adamts9* cKO, *Ift88* cKO, *Tmem67* KO and *Tmem67 ΔCLE* kidneys show transcriptional upregulation of immune activity in the cystic kidneys (n=4 kidneys/ genotype). Error bars indicate Mean ± SEM, **** indicates a p-value <0.0001, ***<0.001, **<0.01, and *<0.05 calculated by two-tailed unpaired *t*-test. Scale bar in **A** is 100 µm, 2 µm in **B,** 50 µm and 30 µm in **C**.

**Figure S8.**
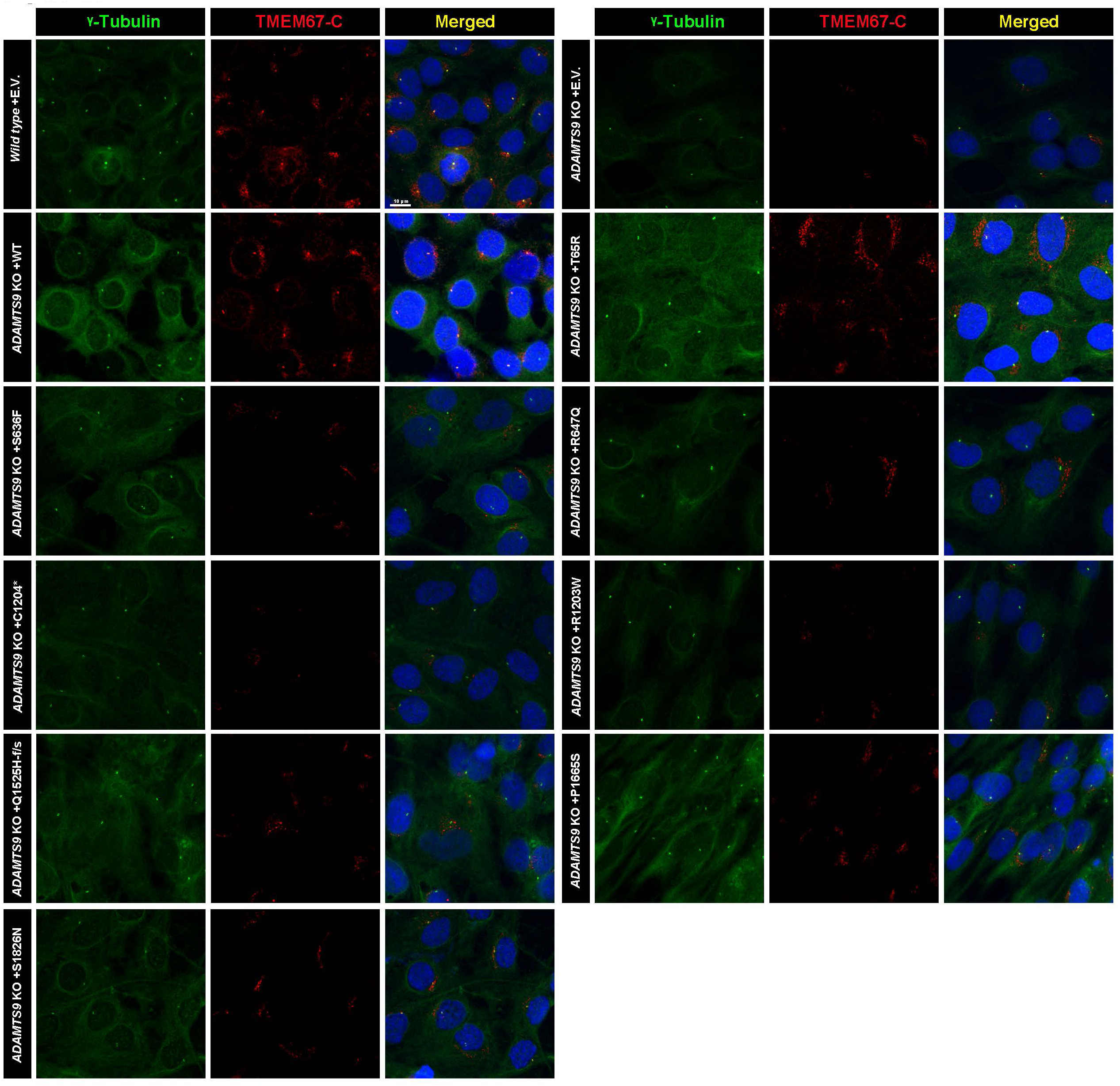
Low magnification images of TMEM67-C immunostaining of ice-cold methanol fixed RPE-1 cells transfected with *ADAMTS9* ciliopathy variants. Low magnification images of immunostaining for TMEM67-C terminus (red) and γ-Tubulin (green) in serum starved and ice-cold methanol fixed cells show significantly reduced transition zone localized TMEM67-C terminus in *ADAMTS9* KO RPE-1 cells, and cells transfected with ciliopathy variants.

**Figure S9.**
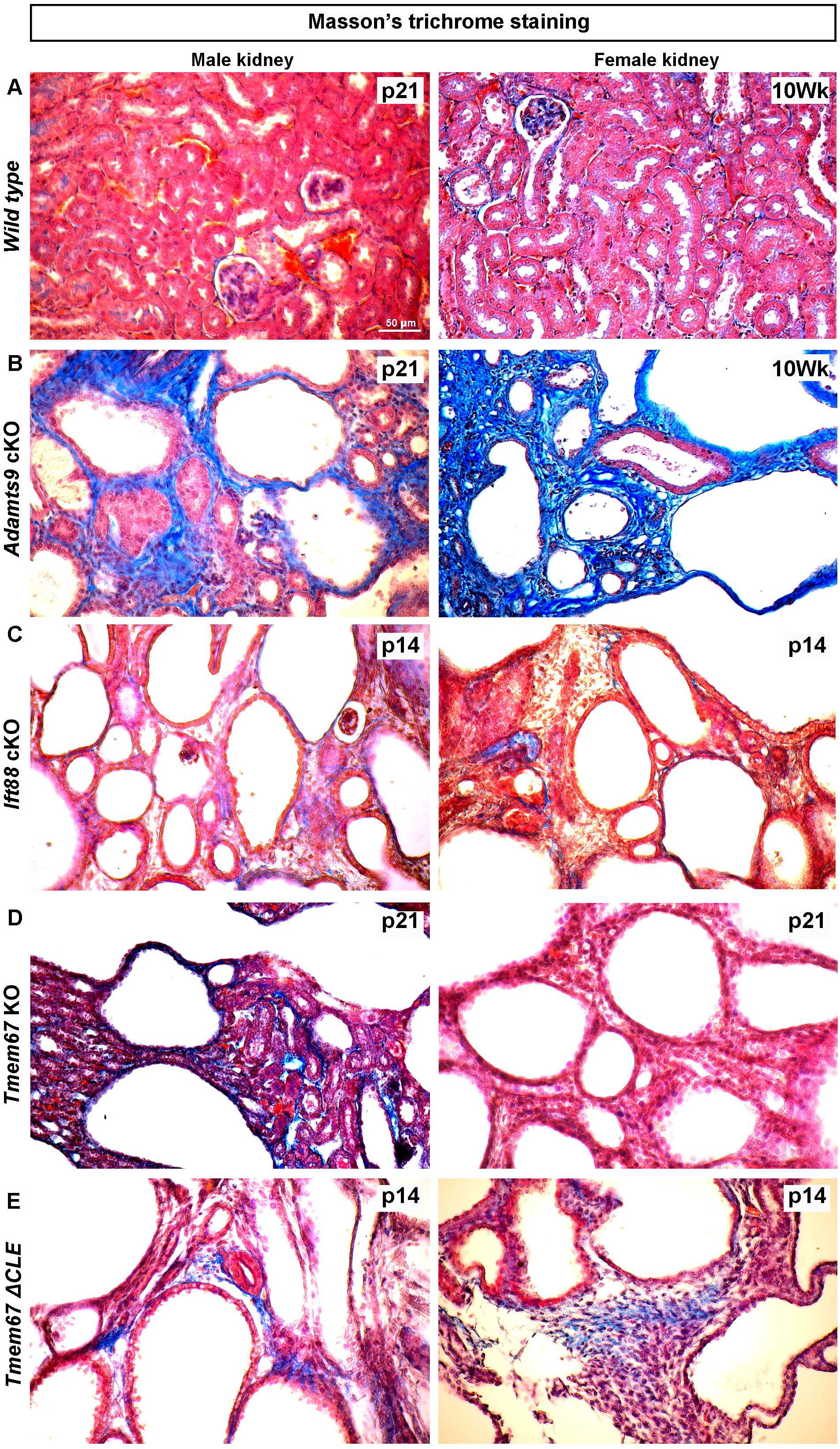
Comparably higher levels of collagen accumulation in *Adamts9* cKO cystic kidneys. (**A-D**) Low magnification images comparing Masson’s trichrome stained kidney sections of *Wt* (**A**), *Adamts9* cKO (**B**), *Ift88* cKO (**C**), *Tmem67* KO (**D**), and *Tmem67 ΔCLE* kidneys (**E**), show distinctly higher levels of collagen (blue staining) present throughout the cystic kidneys of both male (p21) and female (10 week old) *Adamts9* cKO kidneys which may be suppressing the rapid enlargement of *Adamts9* cKO cystic kidneys in comparison to the other cystic kidney models.

**Figure S10.**
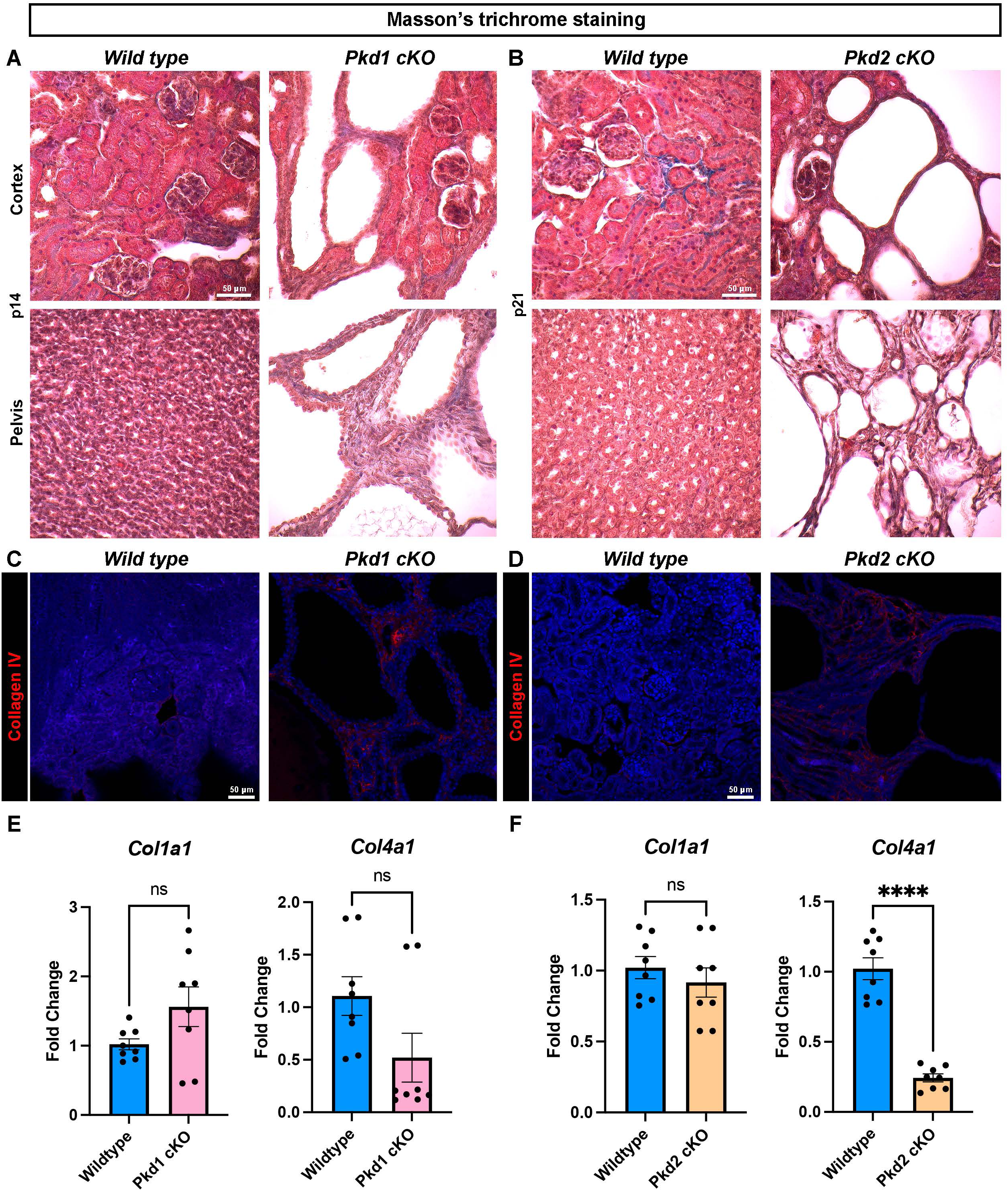
Comparably low levels of collagen in end stage *Pkd1 and Pkd2* cKO cystic kidneys. (**A-B**) Low magnification images comparing Masson’s trichrome stained kidney sections of male *Wt* and *Pkd1* cKO *(Hoxb7-Cre; Pkd1^Fl/Fl^*) kidneys at p14 (**A**), and *Pkd2* cKO *(Rosa26-Cre^ERT^; Pkd2^Fl/Fl^*) kidneys at p21 (**B**), show very minimal levels of collagen (blue staining) present in the *Pkd* cKO cystic kidney models (n=5 male kidneys from each group). (**C-D**) Collagen-IV immunostaining show very low level of little staining in *Pkd1* and *Pkd2* cKO kidneys (n=5 male kidneys from each group). (**E-F**) qRT-PCR analysis shows no significant changes in *Col1a1* transcription while *Col4a1* is down regulated in *Pkd1* and *Pkd2* cKO kidneys (n=4 male kidneys from each group). Error bars indicate Mean ± SEM, **** indicates a p-value <0.0001calculated by two-tailed unpaired *t*-test. Scale bars in **A-D** are 50 µm.

## SUPPLEMENTAL MOVIE LEGENDS

**Supplemental Movie 1 caption:** Movie of 4-week-old male mice showing the distinctly small and hunched posture and abnormal gating of an *Adamts9* cKO mouse and two heterozygous littermates who are phenotypically and behaviorally normal.

**Supplemental Movie 2 caption:** Movie of female littermates of the male mice shown in supplemental movie-1, showing two *Adamts9* cKO and a heterozygous female at 4-weeks of age, who are visually and behaviorally normal and are indistinguishable from one another.

**Table S1:**
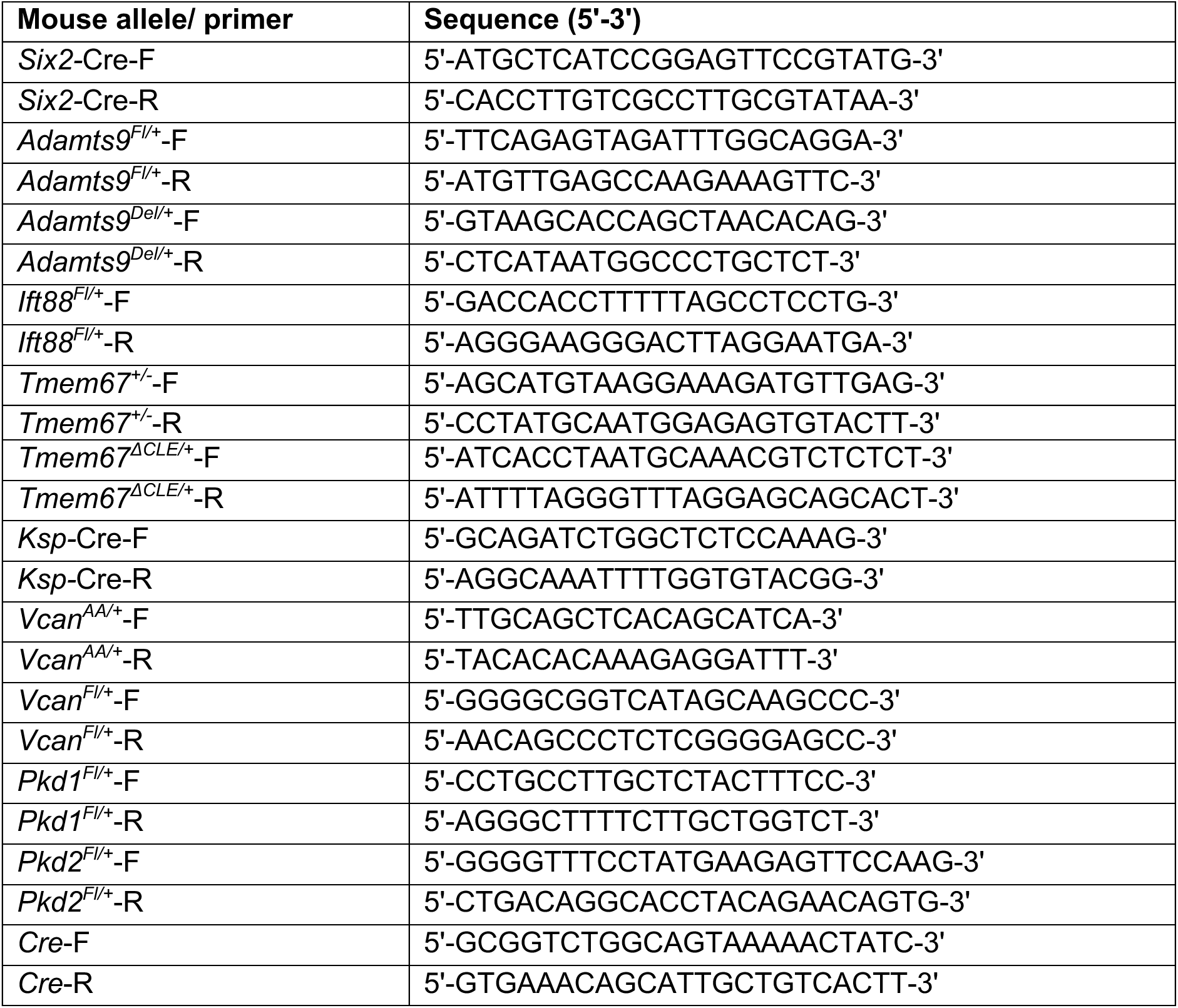
Mouse genotyping primers.

**Table S2:**
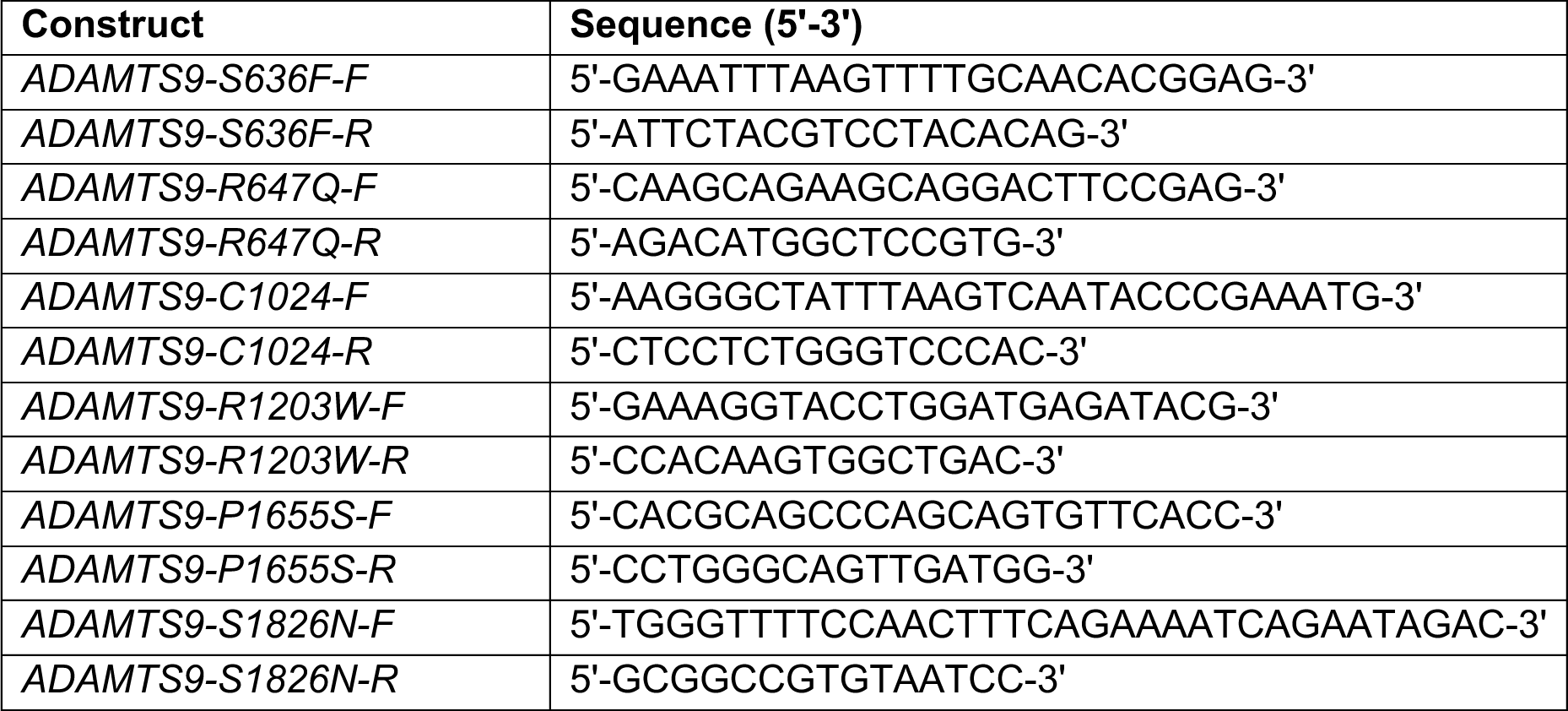
Primers utilized in site directed mutagenesis of *ADAMTS9*.

**Table S3:**
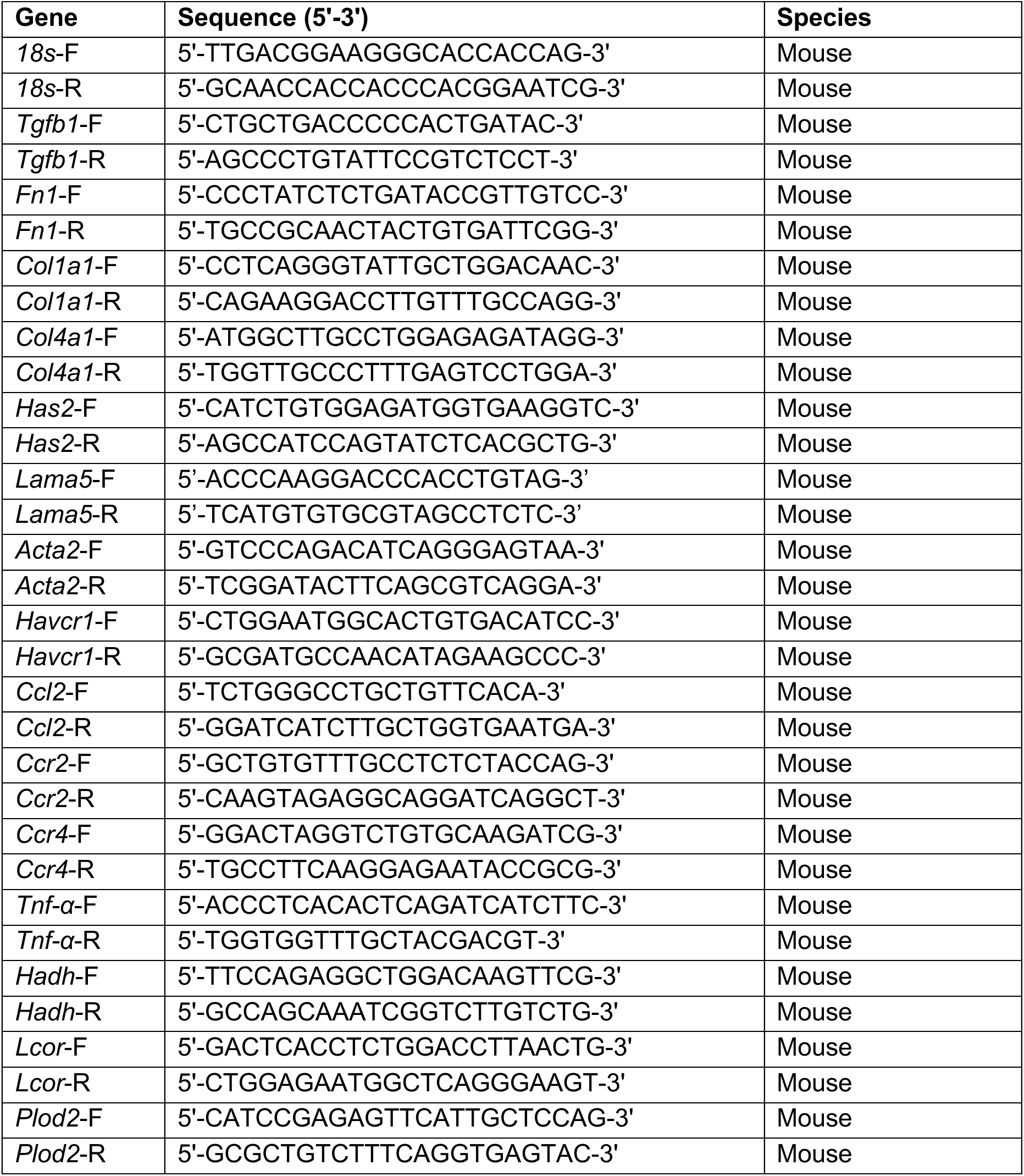
qRT-PCR primers used in this study.

**Table S4:**
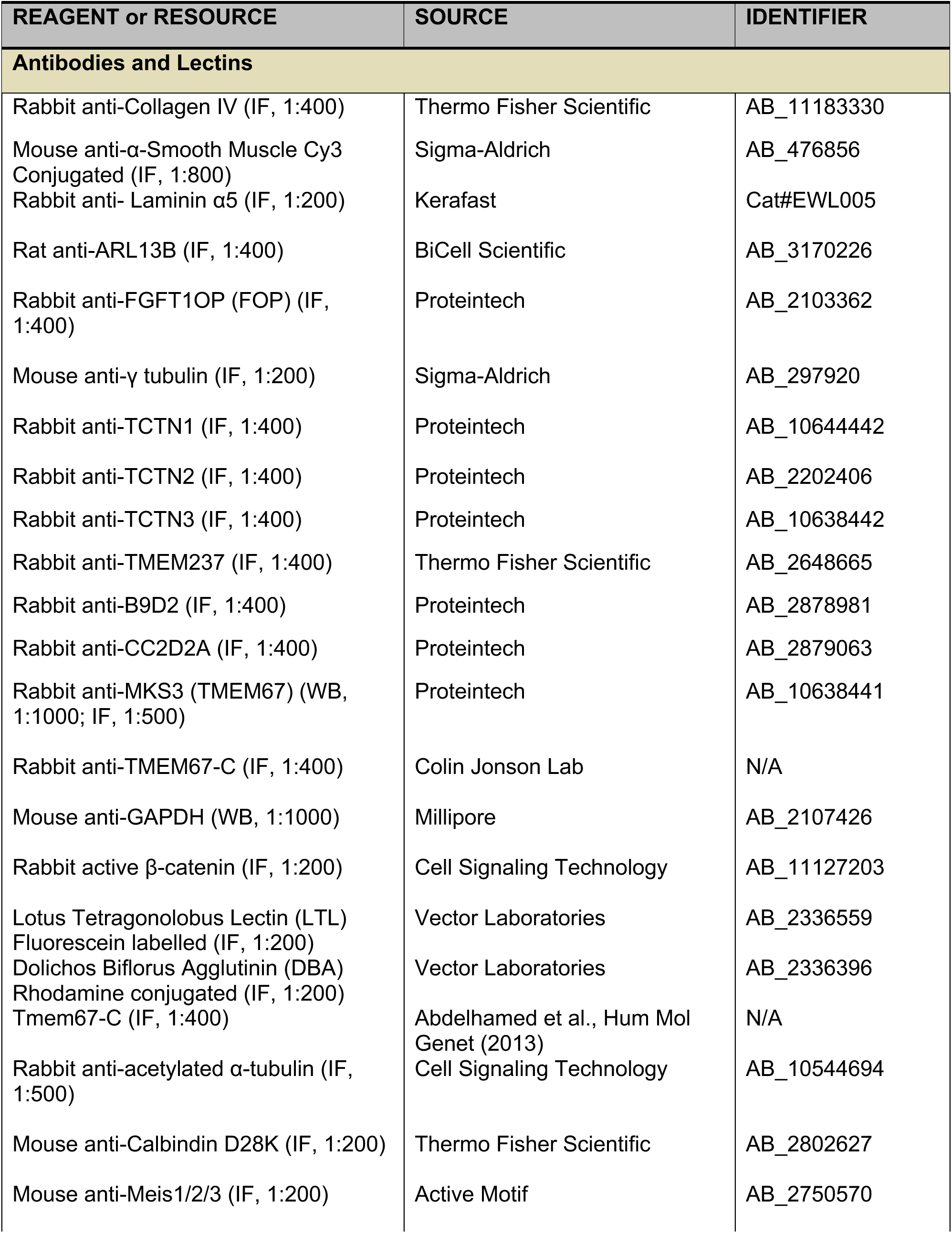
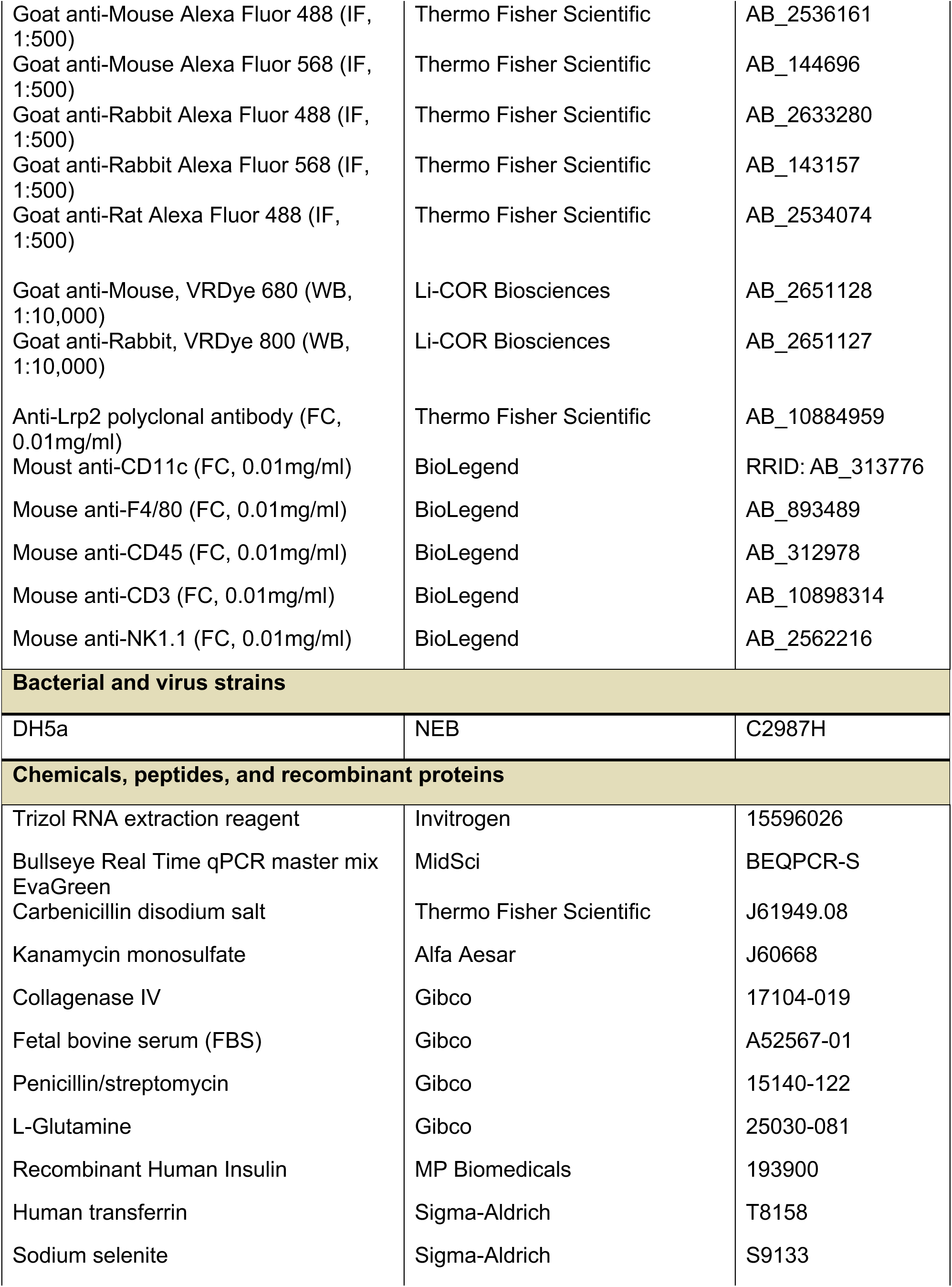
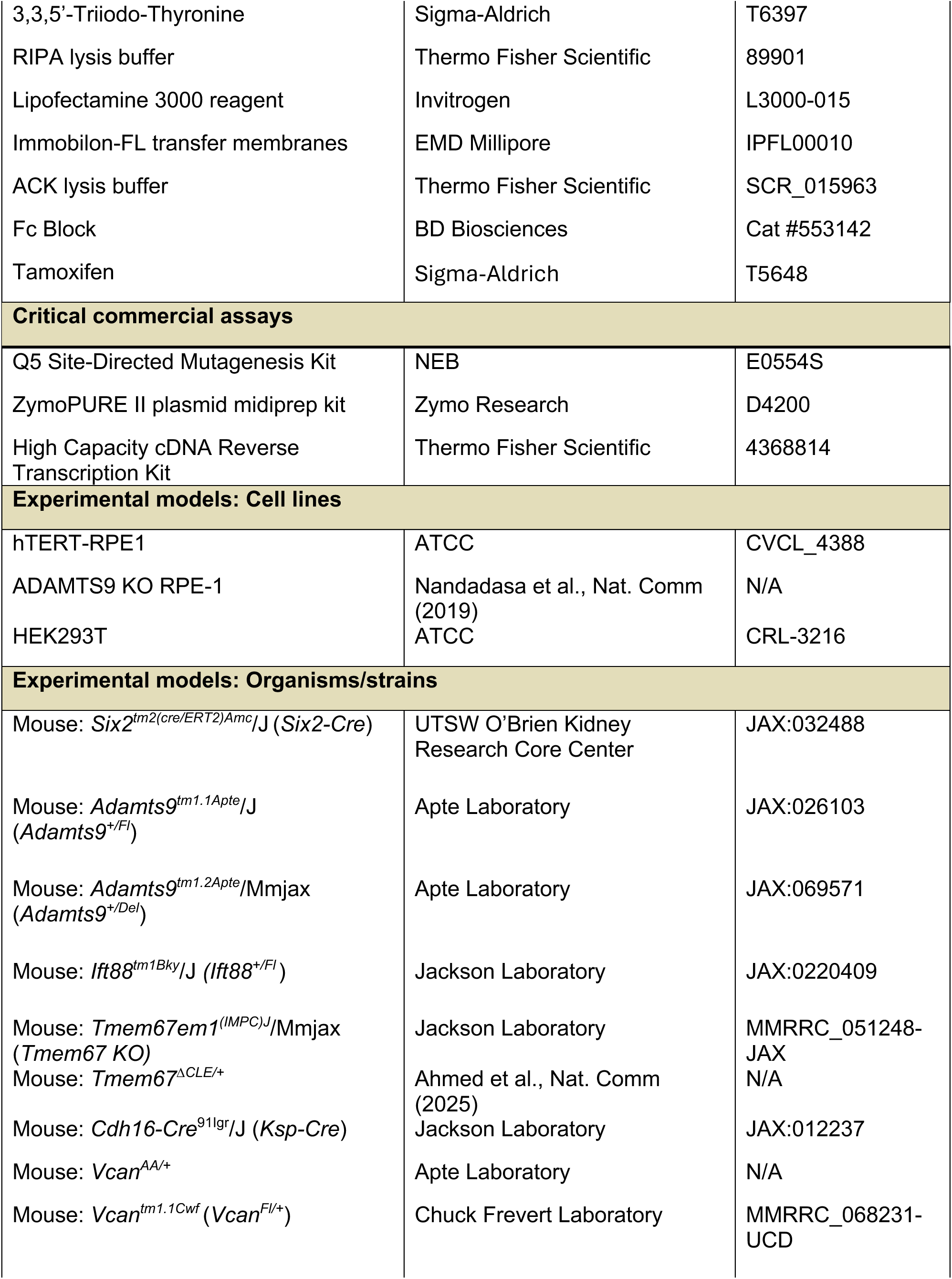
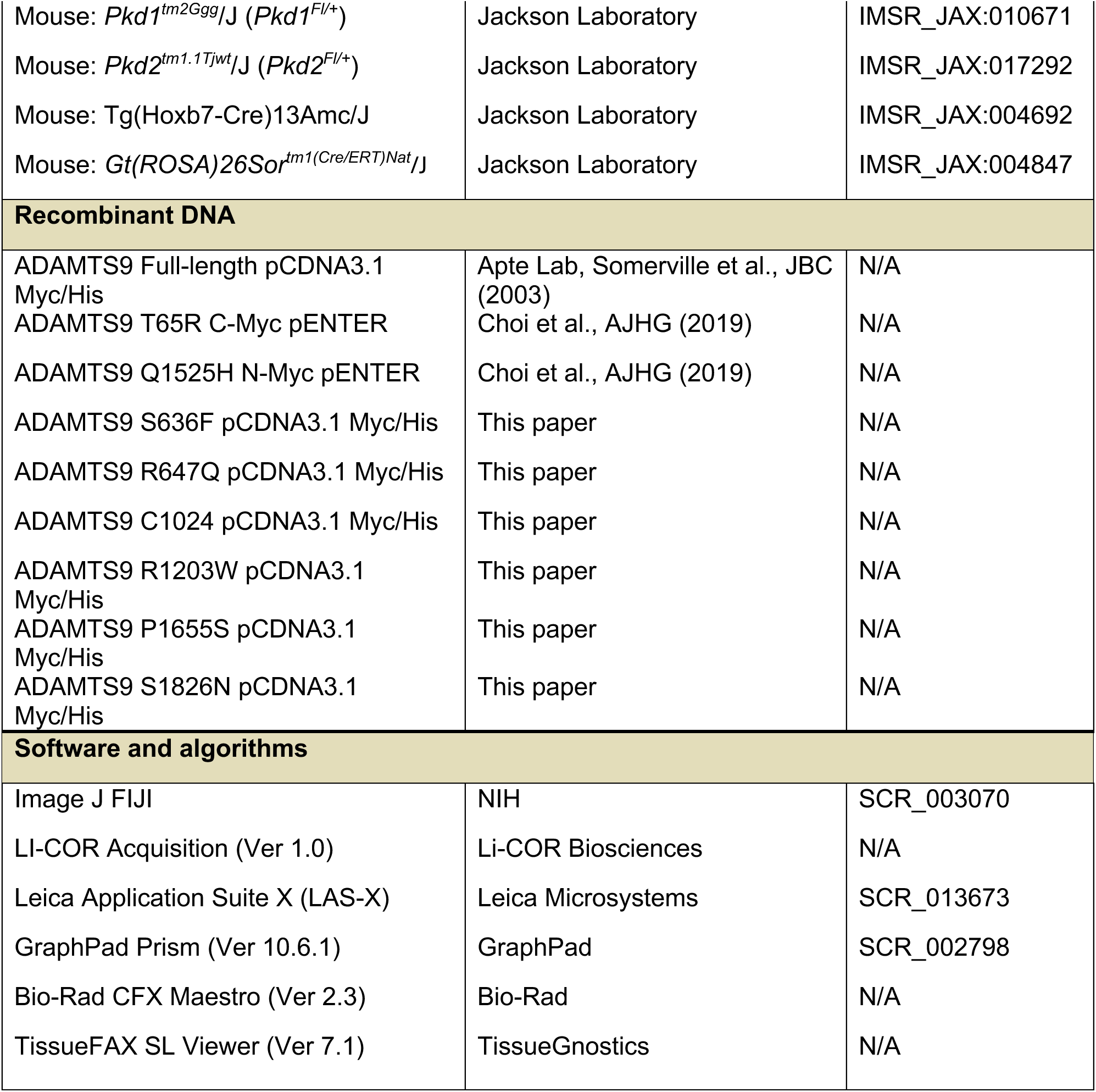
Key Resource Table.

## Notes

### Competing Interest Statement

The authors have declared no competing interest.

### Summary of Updates

Collagen homeostasis in Pkd1 and Pkd2 deleted cystic kidney models were investigated and added to the manuscript as new Figure S10.

